# Thioesterase induction by 2,3,7,8-tetrachlorodibenzo-p-dioxin results in a futile cycle that inhibits hepatic β-oxidation

**DOI:** 10.1101/2021.01.21.427582

**Authors:** Giovan N. Cholico, Russ R. Fling, Nicholas A. Zacharewski, Kelly A. Fader, Rance Nault, Tim Zacharewski

## Abstract

2,3,7,8-Tetrachlorodibenzo-*p*-dioxin (TCDD), a persistent environmental contaminant, induces steatosis by increasing hepatic uptake of dietary and mobilized peripheral fats, inhibiting lipoprotein export, and repressing β-oxidation. In this study, the mechanism of β-oxidation inhibition was investigated by testing the hypothesis that TCDD dose-dependently repressed straight-chain fatty acid oxidation gene expression in mice following oral gavage every 4 days for 28 days. Untargeted metabolomic analysis revealed a dose-dependent decrease in hepatic acyl-CoA levels, while octenoyl-CoA and dicarboxylic acid levels increased. TCDD also dose-dependently repressed the hepatic gene expression associated with triacylglycerol and cholesterol ester hydrolysis, fatty acid binding proteins, fatty acid activation, and 3-ketoacyl-CoA thiolysis while inducing acyl-CoA hydrolysis. Moreover, octenoyl-CoA blocked the hydration of crotonyl-CoA suggesting short chain enoyl-CoA hydratase (ECHS1) activity was inhibited. Collectively, the integration of metabolomics and RNA-seq data suggested TCDD induced a futile cycle of fatty acid activation and acyl-CoA hydrolysis resulting in incomplete β-oxidation, and the accumulation octenoyl-CoA levels that inhibited the activity of short chain enoyl-CoA hydratase (ECHS1).

## INTRODUCTION

Although the liver is the largest internal organ, it is second to adipose tissue in regard to lipid storage capacity. Approximately 20-30% of hepatic fatty acids are derived from the diet, 5-30% from visceral tissues, 15-25% from chylomicron remnants, and 5-30% from *de novo* lipogenesis.^1^ Under normal circumstances, lipid accumulation is coordinated by regulating uptake, utilization, and export (via very low-density lipoproteins [VLDL]) to ensure triacylglycerol (TAG) and cholesterol ester (CE) levels remain low. Excess fatty acids (FAs) are packaged into TAGs and CEs, not only for storage, but to reduce their potential lipotoxicity.^2,3^ Chronic simple, reversible, fat accumulation, or steatosis, may progress to steatohepatitis with fibrosis, as in the case of non-alcoholic fatty liver disease (NAFLD). NAFLD impairs liver function and increases the risk for more complex metabolic diseases such as: metabolic syndrome (MetS), diabetes, cardiovascular disease and hepatocellular carcinoma (HCC). NAFLD alone is estimated to affect ≥35% of the U.S population and has become the second leading reason for requiring a liver transplant.^4^

NAFLD development is described by a ‘multi-hit hypothesis’ in which consecutive insults collectively promote the progression of liver pathologies such as steatosis, inflammation and fibrosis.^5^ Diverse pharmaceuticals and xenobiotics induce hepatic fat accumulation. For example, pesticides and solvents are the most frequently identified chemicals to induce hepatic steatosis while 2,3,7,8-tetrachlorodibenzo-*p*-dioxin (TCDD) and related compounds exhibit the greatest potency.^6,7^ Previous studies have shown that male mice exhibit greater sensitivity to TCDD-elicited reductions in body weight gain compared to females, despite no change in daily food intake.^8–10^ Hepatic micro- and macro-steatosis, immune cell infiltration, fibrosis, and bile duct proliferation (males only) were dose-dependently induced following oral gavage with TCDD every 4 days for 28 days.^11,12^ Lipidomic analysis of hepatic steatosis induced by TCDD and related compounds reported marked increases in FAs, TAGs, phospholipids, and CEs.^13–15^ Fat accumulation has been attributed to increased hepatic uptake of dietary and mobilized peripheral fats, inhibition of VLDL export, and repression of FA oxidation.^8,13,16,17^ In humans, TCDD and related compounds are associated with altered lipid homeostasis, including steatosis with fibrosis and inflammation.^6,18–20^ Epidemiological studies also report elevated serum TAG and cholesterol levels in exposed workers.^21,22^ More recently, a study of second generation Seveso accident victims (age 2-39 years) suggested *in utero* TCDD exposure increased MetS risk in males.^23^ These data suggest that environmental contaminants may play an underappreciated role in the development of NAFLD and its associated metabolic diseases.^24^

Most, if not all, of the effects of TCDD and related compounds are mediated by the aryl hydrocarbon receptor (AHR), a ligand-activated basic-helix-loop-helix Per-Arnt-Sim transcription factor. Although numerous endogenous metabolites, drugs, xenobiotics, and natural products bind to the AHR, its endogenous ligand(s) and role(s) remain elusive.^25^ Ligand binding to the cytosolic AHR causes dissociation of chaperone proteins, followed by translocation to the nucleus and heterodimerization with the aryl hydrocarbon receptor nuclear translocator (ARNT). In the canonical pathway, the AHR-ARNT complex interacts with dioxin response elements (DREs) within the promoter region of target genes, leading to recruitment of transcriptional co-regulators and differential gene expression.^26^ Studies also report binding to non-consensus DREs and DRE-independent mechanisms of differential gene expression.^25,27^ The dose-dependent species-, sex-, age-, tissue-, and cell-specific biochemical and toxicological responses are believed to be the result of aberrant differential gene expression mediated by the AHR.^25,28^ Despite our detailed understanding of AHR-mediated gene regulation, links between TCDD-elicited differential expression and adverse biological effects have not been fully elucidated.

Though FAs can undergo α- and ω-oxidation, β-oxidation is the primary pathway for FA oxidation to produce ATP, as well as intermediates for macromolecular and ketone biosynthesis. Binding proteins, such as diazepam binding inhibitor (DBI; aka acyl-CoA binding protein), fatty acid binding proteins (FABPs), and sterol carrier protein 2 (SCP2), not only shunt FAs to acyl-CoA synthetases for activation, but also protect acyl-CoAs from hydrolysis, channel acyl-CoAs to specific pathways, and protect against free FA and acyl-CoA toxicity.^29–32^ In β-oxidation, activated FAs are subjected to oxidation, hydration, a second oxidation, and finally coenzyme A (CoASH)-dependent thiolytic cleavage to complete the cycle, producing an acetyl-CoA and acyl-CoA that is two carbons shorter.^33^ Peroxisomal and mitochondrial β-oxidation employ similar strategies but exhibit different substrate preferences and use different enzymes encoded by distinct genes. Mitochondrial β-oxidation prefers long, medium, and short chain FAs (LCFAs, MCFAs and SCFAs, respectively), while peroxisomes metabolize very long chain FAs (VLCFAs), LCFAs, bile acid precursor side chains, and dicarboxylic acids (DCAs) produced via ω-oxidation, as well as branched chain FAs following α-oxidation.^34^ Under conditions of acyl-CoA accumulation that limits CoASH availability due to sequestration, cytosolic, peroxisomal, and mitochondrial thioesterases exert control by hydrolyzing β-oxidation substrates and end products, but not intermediates. This releases CoASH in order to support CoASH-dependent reactions in other pathways.^35^

Despite studies reporting β-oxidation inhibition by TCDD and related compounds in rodents, the mechanism remains uncertain. In this study, we examined straight chain FA oxidation to test the hypothesis that TCDD dose-dependently represses gene expression at key steps in β-oxidation. Notably, TCDD decreased hepatic acyl-CoA levels except for the marked dose-dependent increase in octenoyl-CoA and non-monotonic increases in DCAs, consistent with the inhibition of β-oxidation. TCDD repressed genes associated with lipid hydrolysis, FA binding proteins, FA activation, and thiolytic cleavage while inducing thioesterases. These results suggest TCDD induced a futile cycle of FA activation by acyl-CoA synthetases followed by thioesterase-mediated acyl-CoA hydrolysis resulting in incomplete β-oxidation and the accumulation of octenoyl-CoA that inhibited the activity of short chain enoyl-CoA hydratase (ECHS1).

## MATERIALS AND METHODS

### Animal Treatment

Postnatal day 25 (PND25) male C57BL/6 mice weighing within 10% of each other were obtained from Charles River Laboratories (Kingston, NY). Mice were housed in Innovive Innocages (San Diego, CA) containing ALPHA-dri bedding (Shepherd Specialty Papers, Chicago, IL) in a 23°C environment with 30– 40% humidity and a 12 hr/12 hr light/dark cycle. Aquavive water (Innovive) and Harlan Teklad 22/5 Rodent Diet 8940 (Madison, WI) was provided *ad libitum*. On PND28, mice were orally gavaged at the start of the light cycle (zeitgeber [ZT] 0) with 0.1 ml sesame oil vehicle (Sigma-Aldrich, St. Louis, MO) or 0.01, 0.03, 0.1, 0.3, 1, 3, 10, and 30 μg/kg body weight TCDD (AccuStandard, New Haven, CT) every 4 days for 28 days for a total of 7 treatments. The first gavage was administered on day 0, while the final gavage was on day 24 of the 28-day study. This dosing regimen was selected to approach steady state levels given the 8-12 day half-life of TCDD in mice.^36^ Comparable treatment has been used in previous studies.^8,9,13,16,17,37^ On day 28, vehicle- and TCDD-treated mice were weighed and euthanized. Liver samples were immediately flash frozen in liquid nitrogen and stored at -80°C until analysis. All animal procedures were in accordance with the Michigan State University (MSU) Institutional Animal Care and Use Committee ethical guidelines and regulations.

### Liquid Chromatography Tandem Mass Spectrometry

Samples were extracted using methanol:water:chloroform as previously described with slight modifications.^9^ Briefly, untargeted extractions were reconstituted with 400 µl tributylamine with no dilutions for analysis. Analysis was performed using a Xevo G2-XS QTOF attached to a Waters UPLC (Waters, Milford, MA) with negative-mode electrospray ionization run in MS^E^ continuum mode. LC phases, gradient rates, and columns were used as previously published.^9^ For untargeted acyl-CoA analysis, MS^E^ continuum data was processed with Progenesis QI (Waters) to align features, peaks, deconvolute, and identify metabolite peaks. Metabolite identifications were scored based on a mass error <12 ppm to Human metabolome Database entries,^38^ isotopic distribution similarity, and theoretical fragmentation comparisons to MS^E^ high-energy mass spectra using the MetFrag option with each metric contributing a max of 20 points towards a max score of 60. Raw signals for each compound abundances were normalized to a correction factor calculated using a median and mean absolute deviation approach by Progenesis QI. Significance was determined by a one-way ANOVA adjusted for multiple comparisons with a Dunnett’s *post-hoc* test. Raw data is deposited in the NIH Metabolomics Workbench (ST001379).

### Protein Quantification and Capillary Electrophoresis Protein Analysis

Frozen samples (∼50 mg) were homogenized in RIPA buffer with protease inhibitor (Sigma-Aldrich) using a Polytron PT2100 homogenizer (Kinematica, Lucerne, Switzerland) and sonicated on ice. Samples were centrifuged and protein concentration measured using a bovine serum albumin standard curve and a bicinchoninic acid (BCA) assay (Sigma-Aldrich). The WES capillary electrophoresis system (ProteinSimple, San Jose, CA) was used with the following antibodies and dilutions from Proteintech (Rosemont, IL): ACOX1 (1:50; 10957-1-AP), ACSM3 (1:50; 10168-2-AP), ACSL1 (1:50; 13989-1-AP), DBI (1:50; 14490-1-AP). Primary antibodies were detected using an anti-rabbit detection module (ProteinSimple). Chemiluminescence signal intensity was analyzed with Compass Software (ProteinSimple). GraphPad Prism v8.4.3 (La Jolla, CA, USA) was used to conduct a one-way ANOVA followed by a Dunnett’s *post-hoc* analysis to determine statistical significance (p ≤ 0.05).

### ECHS1 Enzymatic Activity Assay

Enoyl-CoA hydratase, short chain 1 (ECHS1) activity was measured spectrophotometrically in hepatic extracts prepared from control mice treated with sesame oil vehicle every 4 days for 28 days using crotonyl-CoA (Sigma-Aldrich) and octenoyl-CoA (Toronto Research Chemicals, Toronto, Canada) as substrates. Total protein lysates were isolated from frozen samples with NP-40 cell lysis buffer (Thermo Fisher Scientific, Waltham, MA) with protease inhibitor using a Polytron PT2100 homogenizer (Kinematica). Incubations were performed in 100mM Tris buffer (pH 8.0) supplemented with 0.1 mg/ml bovine serum albumin at 37°C for 5 min. Reactions were started with 0.02 mg/ml total protein lysate and one of the following substrate ratios: 0.025 mM crotonyl-CoA (1:0 ratio), 0.025 mM crotonyl-CoA:0.025 mM octenoyl-CoA (1:1 ratio), or 0.025 mM crotonyl-CoA:0.25 mM octenoyl-CoA (1:10 ratio). Absorbance was monitored in UV cuvettes at 263 nm using a SpectraMax ABS Plus (Molecular Devices, San Jose, CA). Enzymatic activity was calculated using 6700 as the extinction coefficient.

### Gene Expression, ChIP, pDRE and Protein Location Data

Hepatic RNA-seq data sets were previously published.^9,10,39^ Differentially expressed genes (|fold-change| ≥ 1.5 and posterior probability (P1(*t*)) ≥ 0.8) were determined by empirical Bayes analysis.^40^ Time course (GSE109863), dose response (GSE87519), and diurnal rhythmicity (GSE119780) sequencing data are available at the Gene Expression Omnibus. Diurnal rhythmicity was determined using JTK_CYCLE as previously described.^9^ AHR ChIP-seq (GSE97634) and computationally identified putative dioxin response elements (pDREs, https://doi.org/10.7910/DVN/JASCVZ) data were previously published.^10^ ChIP-seq analysis used a false discovery rate (FDR) ≤ 0.05. pDREs were considered functional with a matrix similarity score (MSS) ≥ 0.856 and associated with genes when located 10kb upstream of the transcription start site (TSS) to the transcription end site (TES). COMPARTMENTS (accessed September 30, 2020) was used for cytosol (C), endoplasmic reticulum (ER), extracellular space (ES), Golgi apparatus (GA), lipid droplet (LD), lysosome (L), mitochondrion (M), mitochondrial outer membrane (OMM), mitochondrial inner membrane (IMM), nucleus (N), peroxisome (P), and plasma membrane (PM) mouse protein localizations.^41^

## RESULTS

### MS analysis of acyl-CoA species

Untargeted metabolomics identified dose-dependent decreases in hepatic acyl-CoAs following oral gavage every 4 days for 28 days (**Table 1**). Octanoyl-, hexanoyl-, butryl- and acetyl-CoA were identified based on parent ion mass, isotope similarity, and theoretical fragmentation. Scores >40 had features matching parent ion mass and isotope distribution and MSE fragmentation data matching *in silico* mass fragmentation, while most metabolite scores averaged ∼35 based on parent ion mass and isotope distribution metrics. The presence of 426.1 m/z and 408.0 m/z coenzyme A fragment ions observed in the MS^E^ fragmentation mass spectra further confirmed the acyl-CoA identifications for octanoyl-, hexanoyl-, butyrl-, and acetyl-CoA.^42^ Hepatic levels of hexanoyl-, butyryl-, and acetyl-CoA were repressed 34.9-, 11.8-, and 6.3-fold at 30 μg/kg TCDD, respectively. Other β-oxidation metabolites also exhibited dose-dependent decreases including butenoyl-CoA, 3-hydroxyhexanoyl-CoA and 3-hydroxybutanoyl-CoA, while octenoyl-CoA was dose-dependently induced 138.9-fold (**Table 1, Supp. Table 1**). These dose-dependent decreases in FA oxidation intermediates are consistent with β-oxidation inhibition.^13,17^

**Table 1:** Effect of TCDD on acyl-CoA levels. Acyl-CoA levels were assessed using untargeted liquid chromatography tandem mass spectrometry. Mice (n=4-5) were orally gavaged every 4 days for 28 days with sesame oil vehicle or TCDD. Bold font and asterisks (^**\***^) denote statistical significance (*p* ≤ 0.05) determined using a one-way ANOVA with a Dunnett’s *post-hoc* analysis. Scores were determined by Progenesis with 60 being the maximum value and 0 being the minimum value. Scores ranging from 30 − 40 are based on mass error and isotope distribution similarity, while score >40 are based on mass error, isotope distribution and fragmentation score. All identified compounds have a score distribution averaging ∼35.

| Compound ID | Description | C<sub>n</sub> | Score | Fold-Change (TCDD vs. Veh.) |  |  |  |  |
| --- | --- | --- | --- | --- | --- | --- | --- | --- |
|  |  |  |  | 0.3 µg/kg | 1 µg/kg | 3 µg/kg | 10 µg/kg | 30 µg/kg |
| HMDB01070 | Octanoyl-CoA | 8 | 36.8 | 1.77 ± 0.63 | 0.04 ± 0.01 | 0.28 ± 0.04 | 0.06 ± 0.03 | 0.01 ± 0.01 |
| HMDB02845 | Hexanoyl-CoA | 6 | 37.2 | 1.43 ± 0.47 | **0.08 ± 0.02\*** | 0.36 ± 0.05 | 0.08 ± 0.02 | **0.03 ± 0.02\*** |
| HMDB01088 | Butyryl-CoA | 4 | 44.3 | 1.24 ± 0.33 | **0.17 ± 0.05\*** | 0.57 ± 0.10 | **0.09 ± 0.01\*** | **0.08 ± 0.04\*** |
| HMDB01206 | Acetyl-CoA | 2 | 52.1 | 0.95 ± 0.12 | **0.30 ± 0.05\*** | **0.53 ± 0.07\*** | **0.04 ± 0.01\*** | **0.16 ± 0.06\*** |
| HMDB03949 | Octenoyl-CoA | 8 | 34 | 1.66 ± 0.19 | 1.14 ± 0.23 | 26.16 ± 25.31 | **114.31 ± 4.89\*** | **139.82 ± 6.88\*** |

### TAG and CE hydrolysis

**Figure 1** summarizes the dose-dependent effects of TCDD on lipases, carboxyesterases, and a deacylase associated with TAG and CE hydrolysis. Note that all fold-changes discussed in the text were derived from the circadian gene expression data set unless otherwise indicated. Despite similar trends, there may be fold-change discrepancies between the dose response and diurnal rhythmicity studies since the former study was not controlled for sample time collection.

**Fig 1:**
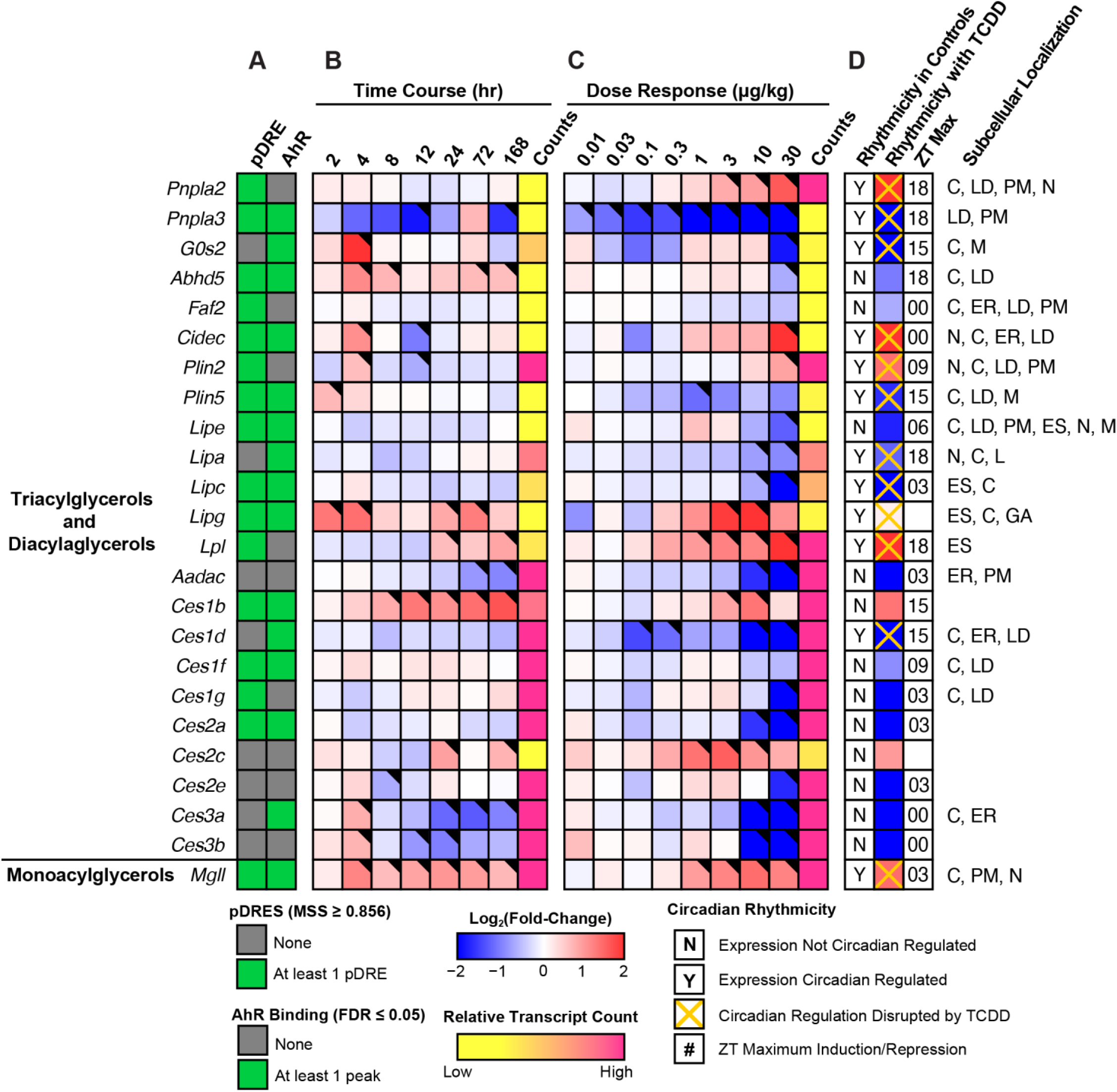
Effect of TCDD on lipid hydrolysis gene expression. Differential expression of genes associated with triacylglycerol hydrolysis assessed using RNA-seq. Mouse Genome Informatics (MGI) official gene symbols are used. **(A)** The presence of putative dioxin response elements (pDREs) and AHR genomic binding at 2 hrs. **(B)** Time-dependent expression following a single bolus dose of 30 μg/kg TCDD (n=3). **(C)** Dose-dependent expression assessed following oral gavage every 4 days for 28 days with TCDD (n=3). **(D)** Diurnal regulated gene expression denoted with a “Y”. An orange ‘X’ indicates abolished diurnal rhythm following oral gavage with 30 μg/kg TCDD every 4 days for 28 days. ZT indicates statistically significant (P1(*t*) > 0.8) time of maximum induction/repression. Counts represents the maximum number of raw aligned reads for any treatment group. Low counts (<500 reads) are denoted in yellow with high counts (>10,000) in pink. Differential gene expression with aposterior probability (P1(*t*)) >0.80 is indicated by a black triangle in the top right tile corner. Protein subcellular locations were obtained from COMPARTMENTS and abbreviated as: cytosol (C), endoplasmic reticulum (ER), extracellular space (ES), Golgi apparatus (GA), lipid droplet (LD), lysosome (L), mitochondrion (M), nucleus (N), and plasma membrane (PM).

Although TCDD induced gene expression of adipose triglyceride lipase (ATGL, *Pnpla2*) 4.3-fold, its paralog, *Pnpla3*, was repressed 37.7-fold. The requisite ATGL co-activator, CGI-58 (Abhydrolase Domain Containing 5, *Abhd5*) was also repressed 2.0-fold, as was *G0s2* (10.9-fold), a potent ATGL inhibitor.^43^ In addition, other ATGL regulators including *Faf2* (−1.6-fold), *Cidec* (8.7-fold), *Plin2* (2.6-fold) and *Plin5* (−3.1-fold) exhibited differential expression.^44^ Lysosomal lipase A (*Lipa*, 2.3-fold), hormone sensitive lipase (*Lipe*, 3.3-fold), and hepatic lipase C (*Lipc*, 12.0-fold), which deacylate endocytosed low-density lipoprotein TAGs and CEs, were dose-dependently repressed. Highly expressed carboxyesterases, which serve important roles in lipid metabolism and VLDL assembly,^45^ exhibited dose-dependent repression with the exception of *Ces1b*. Specifically, highly expressed *Ces2a, Ces2e, Ces3a*, and *Ces3b* were repressed 56.3-, 4.0-, 1,923-, and 1,333-fold, respectively. *Ces* mRNA, protein and/or enzymatic activity repression by TCDD and related compounds in mice and rats has been previously reported.^37,46,47^ Arylamide deacetylase (*Aadac*), another highly expressed gene associated with hepatic TAG metabolism,^43^ was dose-dependently repressed 28.6-fold. Both CESs and AADAC are localized to the extracellular region or endoplasmic reticulum. Interestingly, ATGL and monoglyceride lipase (*Mgll*, 2.7-fold), which are primarily located in the nucleus, cytosol, and plasma membranes, were both induced. In addition to substrate preferences based on fatty acid composition and cellular location, hydrolyases channel released FAs to specific fates such as β-oxidation, membrane formation, VLDL assembly or PPAR activation.^43,45^ Some lipases and carboxyesterases also exhibit cholesterol and retinyl ester hydrolysis activity.

The differential expression of genes associated with lipid hydrolysis was more pronounced at 28 days in mice treated with 30 μg/kg TCDD. Differential expression did not occur exclusively with genes exhibiting AHR enrichment. Although TCDD disrupted the diurnal rhythmicity of most lipid hydrolysis genes, this was not the case for highly expressed carboxyesterases which did not exhibit diurnal regulation. *Ces* genes are localized to a tandem cluster on chromosome 8 which may explain their collective repression by TCDD except for *Ces1b*.^45^ *Ces* repression may also be due to the induction of proinflammatory cytokine signaling.^37^ Overall, the effect of TCDD on gene expression suggests lipid hydrolysis was repressed. However, cellular lipase mRNA levels do not always correlate with enzyme activity due to extensive post-translational regulation.^48^ Accordingly, hepatic levels of TAG, CEs, and bile acids were higher, as were free FAs suggesting esterification may be saturated.^13,15,41^

### FA and acyl-CoA binding proteins

We next examined the effect of TCDD on binding proteins that are important for lipid uptake, intra-/extra-cellular trafficking, and cytoprotection (**Fig. 2**).^31,32^ In addition to channeling lipids to specific metabolic pathways, fatty acid binding proteins (FABPs), acyl-CoA binding protein (ACBP; aka diazepam binding inhibitor [DBI]), and sterol carrier protein 2 (SCP2) mitigate the toxicity, hydrolysis, and signaling potential of free acyl-CoAs. FABPs, DBI, and SCP2 also bind acyl-CoAs as well as other hydrophobic ligands including peroxisome proliferators, prostaglandins, bile acids, bilirubin, heme, fatty acid, and lipid metabolites. Highly expressed *Fabp1, Dbi*, and *Scp2* encode for the majority of acyl-CoA buffering capacity and were repressed 5.9-, 7.1-, and 3.5-fold, respectively. Only *Dbi* exhibited an oscillating expression pattern that was abolished by TCDD. Despite AHR enrichment at 2 hrs in the presence of a putative DRE (pDRE), *Fabp1, Dbi*, and *Scp2* exhibited minimal repression within 168 hrs of a single bolus gavage of 30 µg/kg TCDD. At 28 days, dose-dependent repression of *Fabp1, Dbi*, and *Scp2* was observed with 10 and 30 µg/kg TCDD. *Fabp2* and *5* were also repressed 4.5- and 1.9-fold, respectively. Despite the 6.6-, 14.0-, and 120.1-fold induction of *Fabp4, 7*, and *12*, respectively, their induction would likely be insufficient to compensate for the loss of buffering capacity provided by FABP1, which showed expression levels ∼125-fold higher than the other FABPs. FABP1, unlike other FABPs that only bind one FA, binds two FAs and accounts for 7-11% of the cytosolic protein in normal human liver.^31^ *Fabp1, 5*, and *12* all had multiple AHR enrichment sites but only *Fabp2* and *12* exhibited differential expression at 168 hrs. Acyl-CoA binding domain containing proteins 4 and 5 (*Acbd4* and *5*), which facilitate acyl-CoA transfer to peroxisomes, were also repressed 2.7- and 3.1-fold, respectively. Consequently, decreased binding protein levels may impair FA and acyl-CoA channeling to specific pathways while increasing the potential for toxicity, signaling, and membrane disruption, as well as acyl-CoA susceptibility to hydrolysis.

**Fig 2:**
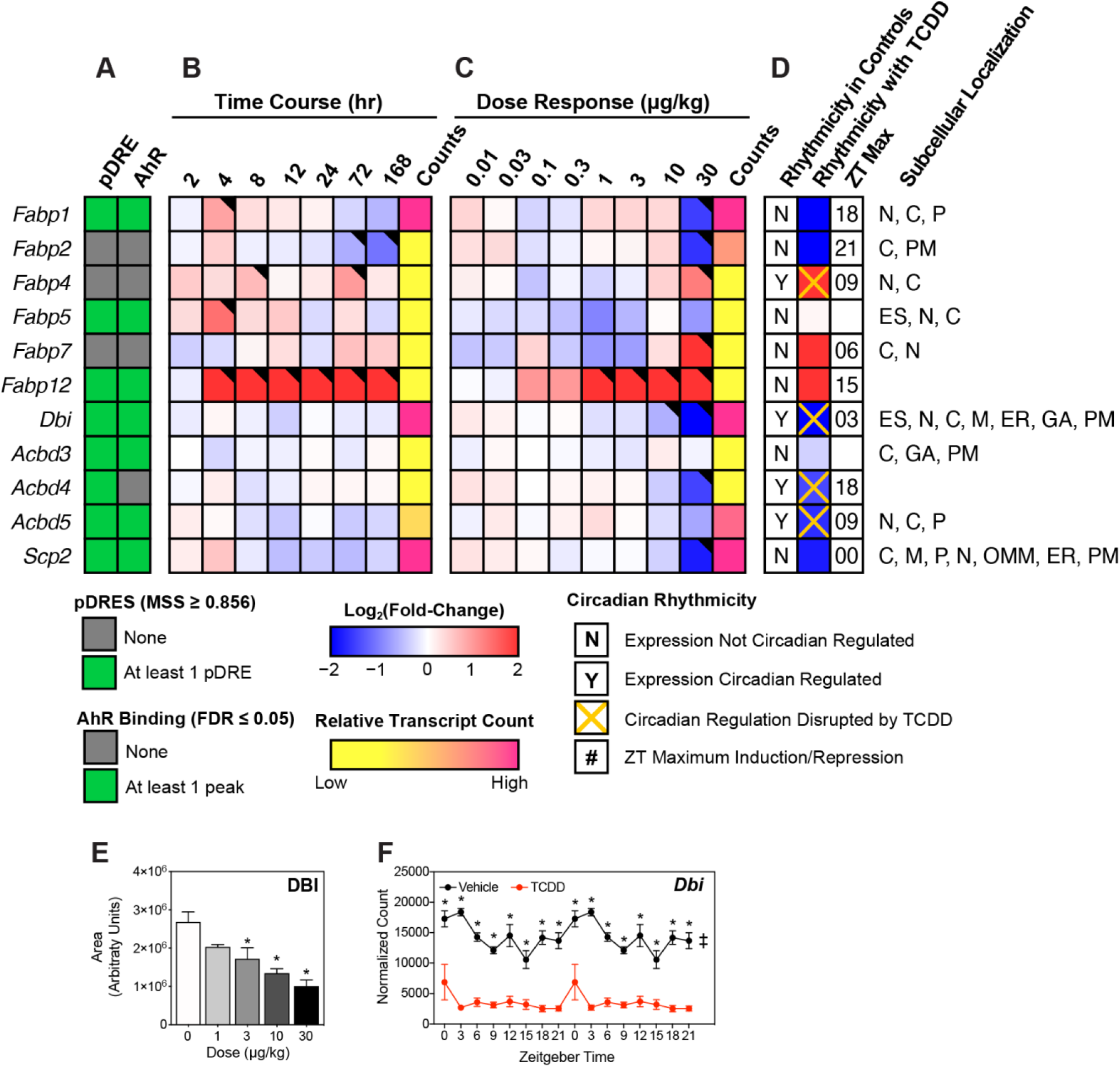
Effect of TCDD on intracellular fatty acid trafficking proteins. Differential expression of genes associated with intracellular fatty acid trafficking assessed using RNA-seq. Mouse Genome Informatics (MGI) official gene symbols are used. **(A)** The presence of putative dioxin response elements (pDREs) and AHR genomic binding at 2 hrs. (**B)** Time-dependent gene expression following a single bolus dose of 30 μg/kg TCDD (n=3). **(C)** Dose-dependent gene expression assessed following oral gavage every 4 days for 28 days with TCDD (n=3). **(D)** Diurnal regulated gene expression denoted with a “Y”. An orange ‘X’ indicates abolished diurnal rhythm following oral gavage with 30 μg/kg TCDD every 4 days for 28 days. ZT indicates statistically significant (P1(*t*) > 0.8) time of maximum gene induction/repression. Counts represent the maximum number of raw aligned reads for any treatment group. Low counts (<500 reads) are denoted in yellow with high counts (>10,000) in pink. Differential gene expression with a posterior probability (P1(*t*)) >0.80 is indicated by a black triangle in the top right tile corner. Protein subcellular locations were obtained from COMPARTMENTS and abbreviated as: cytosol (C), endoplasmic reticulum (ER), extracellular space (ES), Golgi apparatus (GA), mitochondrion (M), outer mitochondrial membrane (OMM), nucleus (N), peroxisome (P), and plasma membrane (PM). **(E)** Capillary electrophoresis was used to assess DBI protein levels in total lysate prepared from liver samples harvested between ZT0-3 (n=3). Bar graphs denote the mean ± SEM. Statistical significance (^*^p ≤ 0.05) was determined using a one-way ANOVA followed by Dunnett’s *post-hoc* analysis. **(F)** Diurnal expression of *Dbi* was assessed by RNA-seq (n=3). Asterisks denotes a posterior probability (P1(*t*) > 0.8) within the same timepoint comparing vehicle to TCDD. Diurnal rhythmicity (‡) was determined using JTK_CYCLE for each treatment. Circadian data are plotted twice along the x-axis to better visualize the gene expression rhythmicity.

### FA activation

Twenty-five mouse acyl-CoA synthetases (ACSs) catalyze the irreversible activation of FAs for β-oxidation or lipid biosynthesis. Each exhibits different tissue expression, subcellular locations, and substrate preferences while also channeling substrates to different pathways.^49^ Twenty-two ACS genes were detected in the mouse liver including short-(*Acss1-2*; carbons length (C) 2-4), medium-(*Acsm1-5*; C6-10), long-(*Acsl1, 3-6*; C12-18), and very long-(*Slc27a1-6*; C12-30) members. Of the 17 abundantly expressed ACS genes, 15 were repressed including highly expressed (>10,000 counts) *Acsm1* (−12.5-fold), *Acsl1* (−9.1-fold), *Acsl5* (−5.6-fold), *Slc27a2* (−5.9-fold), and *Slc27a5* (−16.9-fold) following TCDD treatment (**Fig. 3**). In general, repression was dose-dependently observed within 168 hrs of a single bolus gavage of 30 µg/kg TCDD. Most ACS loci exhibited AHR enrichment in the presence of a pDRE at 2 hrs and exhibited disrupted diurnal expression. Although ACS mRNA levels poorly correlate with protein and enzyme activity,^50^ TCDD-elicited gene repression was consistent with decreases in ACSM3 and ACSL1 protein levels (**Figs. 3F** and **H**). Collectively, these data point to ACS repression by TCDD being a direct target of AHR activation.

**Fig 3:**
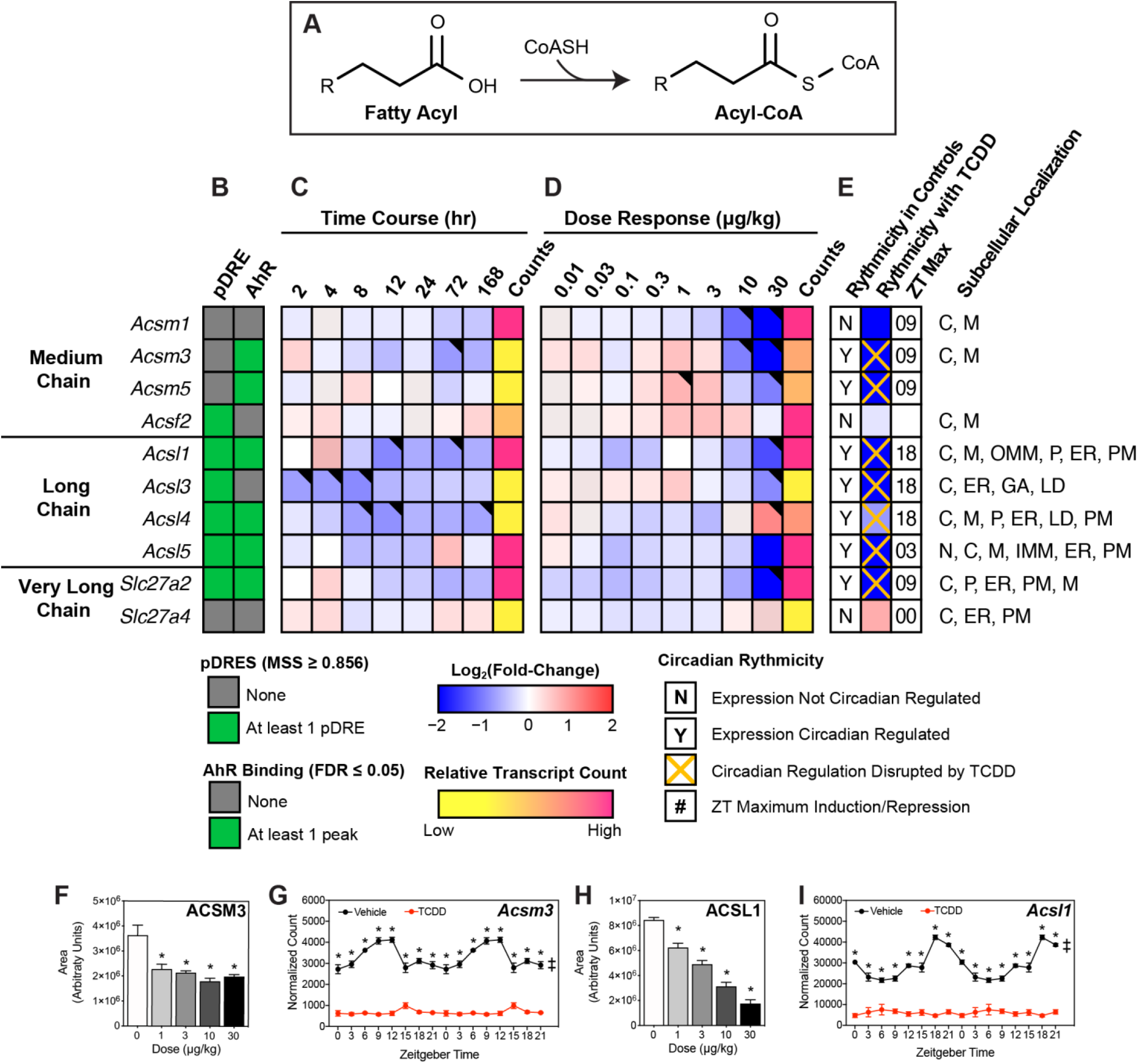
Effect of TCDD on hepatic fatty acid activation. Differential expression of genes associated with fatty acid activation assessed using RNA-seq. **(A)** Fatty acid activation reaction catalyzed via acyl-CoA synthetases. Official gene symbol designated in the MGI database are listed. **(B)** The presence of putative dioxin response elements (pDREs) and AhR enrichment at 2 hrs. (**C)** Time-dependent gene expression was assessed following a single bolus dose of 30 μg/kg TCDD (n=3). **(D)** Dose-dependent gene expression following oral gavaged every 4 days for 28 days with 0.01, 0.03, 0.1, 0.3, 1, 3, 10 or 30 μg/kg TCDD (n=3). **(E)** Circadian regulated genes are denoted with a “Y”. An orange ‘X’ indicates abolished diurnal rhythm following oral gavage with 30 μg/kg TCDD every 4 days for 28 days. ZT indicates statistically significant (P1(*t*) > 0.8) time of maximum gene induction/repression. Counts represent the maximum number of raw aligned reads for any treatment group. Low counts (<500 reads) are denoted in yellow with high counts (>10,000) in pink. Differential expression with a posterior probability (P1(*t*)) >0.80 is indicated with a black triangle in the top right tile corner. Protein subcellular locations were obtained from COMPARTMENTS and abbreviated as: cytosol (C), endoplasmic reticulum (ER), Golgi apparatus (GA), lipid droplet (LD), lysosome (L), mitochondrion (M), outer mitochondrial membrane (OMM), inner mitochondrial membrane (IMM), nucleus (N), peroxisome (P), and plasma membrane (PM). Capillary electrophoresis was used to assess **(F)** ACSM3 and **(H)** ACSL1 protein levels in total lysate prepared from liver samples harvested between ZT0-3 (n=3). Bar graphs denote the mean ± SEM. Statistical significance (^*^p ≤ 0.05) was determined using a one-way ANOVA followed by Dunnett’s post-hoc analysis. Diurnal expression of **(G)** *Acsm3* and **(I)** *Acsl1* was assessed by RNA-seq (n=3). Asterisks denotes a posterior probability (P1(*t*) > 0.8) within the same timepoint comparing vehicle to TCDD. Diurnal rhythmicity (‡) was determined using JTK_CYCLE for each treatment. Circadian data are plotted twice along the x-axis to better visualize the gene expression rhythmicity.

### Mitochondrial and peroxisomal transport

While SCFAs and MCFAs passively enter mitochondria, LCFAs and VLCFAS must be transported into the mitochondria.^51^ At the outer mitochondrial membrane, carnitine palmitoyltransferase I (*Cpt1a*; −2.7-fold) replaces CoASH in the activated FA with carnitine for transport of the acyl-carnitine (**Fig. 4**). Carnitine/acylcarnitine translocase (*Slc25a20* aka CACT; −1.6-fold), transports acyl-carnitine species across the inner mitochondrial membrane into the matrix where carnitine palmitoyltransferase 2 (*Cpt2*; no change in gene expression) reactivates the acyl-carnitine into acyl-CoA in preparation for β-oxidation. *Cpt1a* and *Slc25a20*, which both possess pDREs, were also repressed within 168 hrs with evidence of AHR enrichment at 2 hrs.

**Fig 4:**
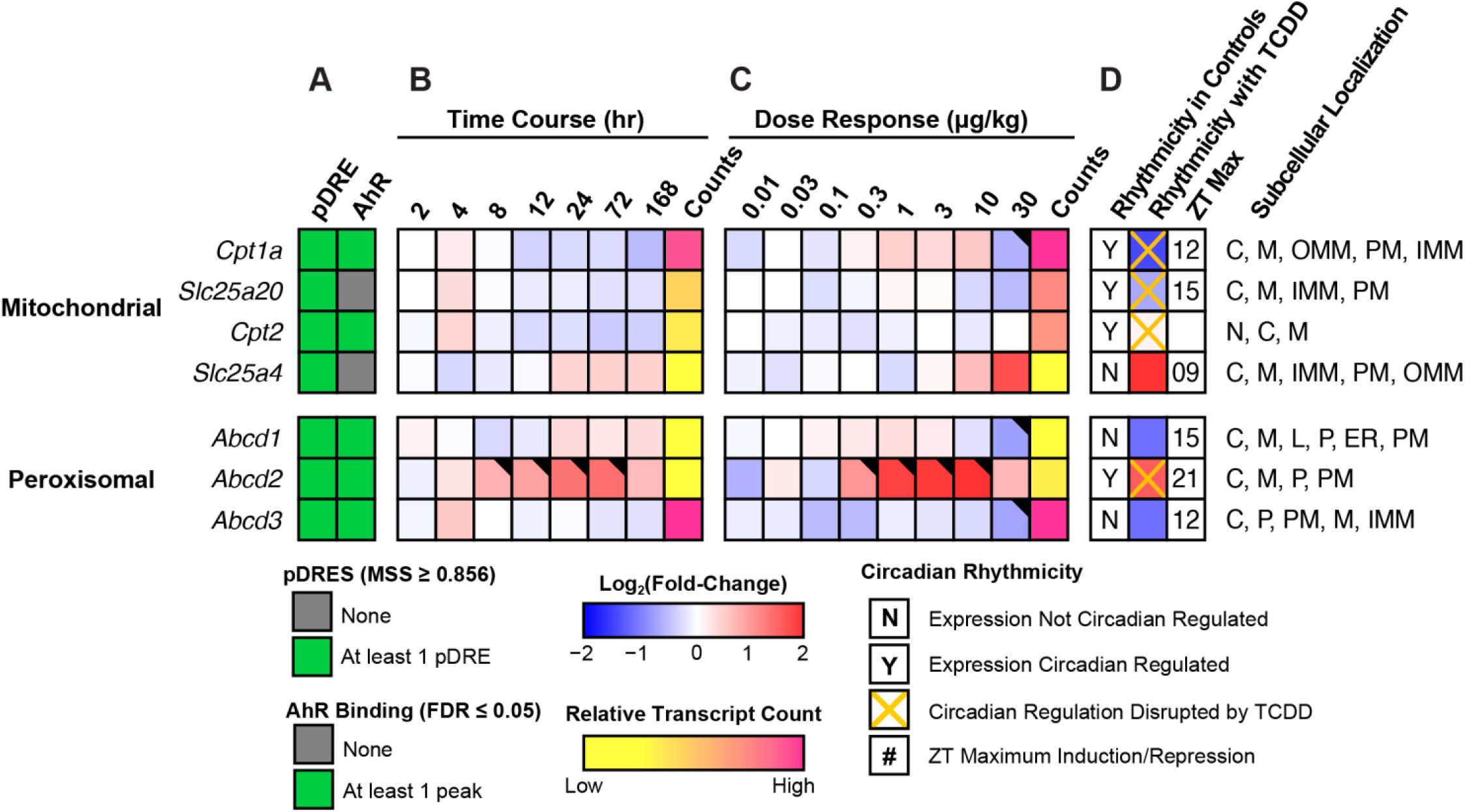
Mitochondrial and peroxisomal free fatty acid and acyl-CoA transport. Differential expression of genes associated with fatty acid transport assessed by RNA-seq. Mouse Genome Informatics (MGI) official gene symbols are used. **(A)** The presence of putative dioxin response elements (pDREs) and AHR genomic binding at 2 hrs. (**B)** Time-dependent expression following a single bolus dose of 30 μg/kg TCDD (n=3). **(C)** Dose-dependent gene expression assessed following oral gavage every 4 days for 28 days with TCDD (n=3). **(D)** Diurnal regulated gene expression denoted with a “Y”. An orange ‘X’ indicates abolished diurnal rhythm following oral gavage with 30 μg/kg TCDD every 4 days for 28 days. ZT indicates statistically significant (P1(*t*) > 0.8) time of maximum gene induction/repression. Counts represent the maximum number of raw aligned reads for any treatment group. Low counts (<500 reads) are denoted in yellow with high counts (>10,000) in pink. Differential gene expression with a posterior probability (P1(*t*)) >0.80 is indicated by a black triangle in the top right tile corner. Protein subcellular locations were obtained from COMPARTMENTS and abbreviated as: cytosol (C), endoplasmic reticulum (ER), lysosome (L), mitochondrion (M), outer mitochondrial membrane (OMM), inner mitochondrial membrane (IMM), nucleus (N), peroxisome (P), and plasma membrane (PM).

ATP binding cassette subfamily member D (ABCD) 1 and 2, and to a lesser extent ABCD3, transport long- and very long-chain acyl-CoAs into peroxisomes.^52^ ABCD1 (*Abcd1*), which prefers saturated and unsaturated 18-22C acyl-CoA substrates, and ABCD3 (*Abcd3*), with a preference for bile acid conjugated FAs,^53^ were both repressed 2.2-fold (**Fig. 4**). *Abcd1* and *3* exhibited negligible differential expression in the time course study despite AHR enrichment with pDREs. In contrast, *Abcd2* showed disrupted oscillating expression in the time course and dose response studies. Although *Abcd2* prefers C22:6- and C24:6-CoAs,^53^ the level of very long unsaturated FAs in TAGs and CEs was increased by TCDD.^13^

### Dehydrogenation of activated acyl-CoA Species

Despite the repression of lipid hydrolases and decreased acyl-CoA binding capacity, **Table 1** indicates ongoing β-oxidation in the presence of TCDD as evidenced by the presence of short- and medium-chain acyl-CoAs. The first step involves acyl-CoA dehydrogenation between C2 and C3 to produce trans-2-enoyl-CoA. In the mitochondria, this oxidation is catalyzed by acyl-CoA dehydrogenases (ACAD) which exhibit different tissue expression, subcellular locations, and substrate preferences. Highly expressed *Acad11*, which preferentially oxidizes saturated C22-CoA, was repressed in the time course and dose response (3.8-fold at 30 µg/kg TCDD) studies with AHR enrichment at 2 hrs in the presence of a pDRE (**Fig. 5**). However, highly expressed *Acadvl* (induced 1.6-fold), which has overlapping substrate preferences, likely offsets the decreased expression of *Acad11* while the effects on *Acadm*, which preferentially oxidizes medium-chain acyl-CoAs, were negligible. Overall, the effects of TCDD on mitochondrial acyl-CoA dehydrogenase gene expression were modest. In contrast, highly expressed acyl-CoA oxidase 1 (ACOX1), the rate-limiting step in peroxisomal β-oxidation, was repressed in the time course, dose response and circadian (3.9-fold at 30 µg/kg TCDD) studies in the presence of AHR enrichment at 2 hrs and pDREs (**Fig. 5**). Accordingly, ACOX1 protein levels were also repressed (**Fig. 5F**). The results suggests peroxisomal β-oxidation is impeded by TCDD, consistent with the accumulation of free and esterified very long- and long-chain FAs within TAGs, CEs, and phospholipids.^13,15^

**Fig 5:**
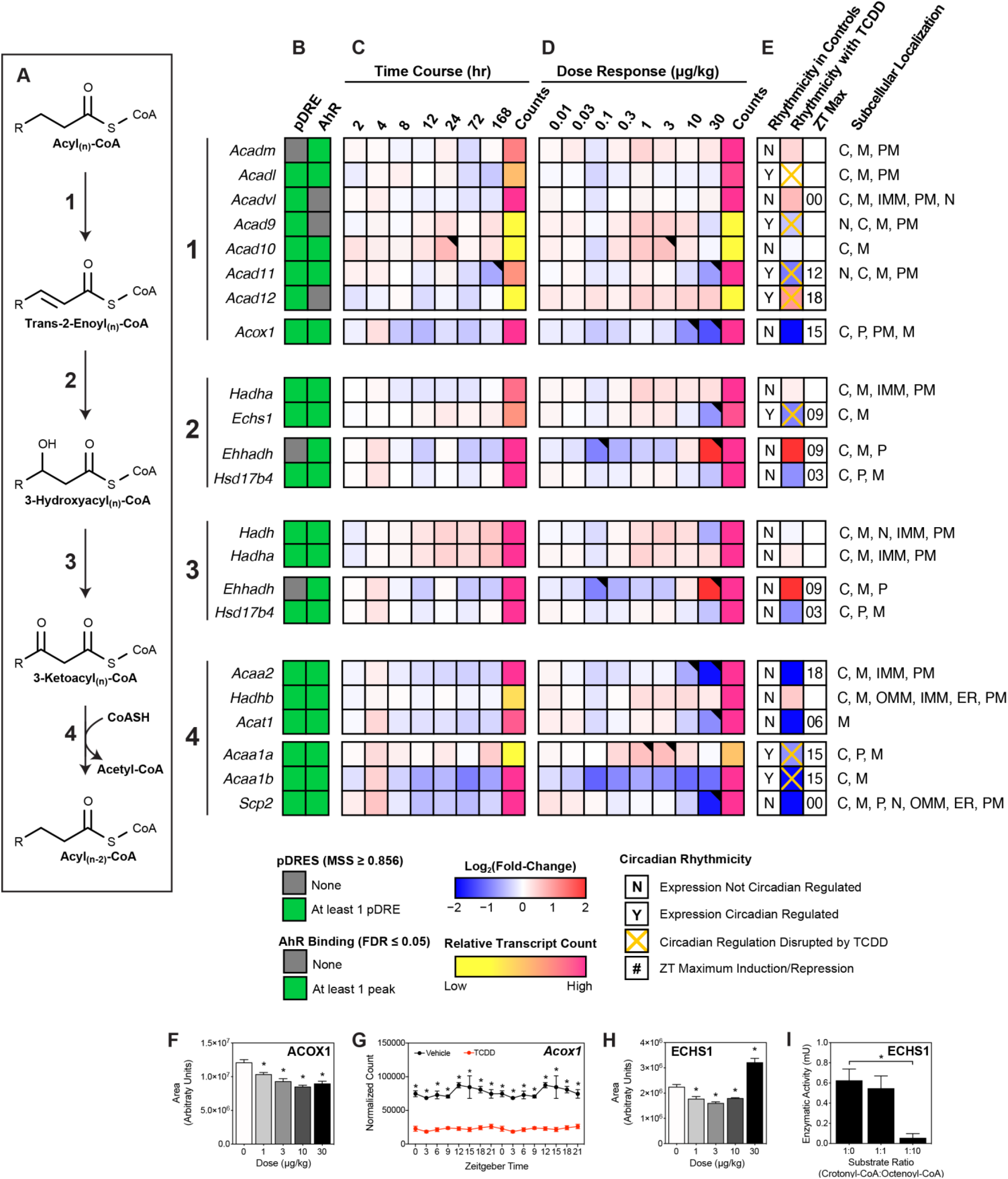
Effect of TCDD on β-Oxidation of acyl-CoAs. Differential expression of genes associated with β-oxidation assessed by RNA-seq. **(A)** Fatty acid β-oxidation is sequentially catalyzed via acyl-CoA dehydrogenases, enoyl-CoA hydratases, hydroxyacyl-CoA dehydrogenases and thiolases. Official gene symbol designated in the MGI database are listed. **(B)** The presence of putative dioxin response elements (pDREs) and AHR enrichment at 2 hrs. **(C)** Time-dependent gene expression was assessed following a single bolus dose of 30 μg/kg TCDD (n=3). **(D)** Dose-dependent gene expression following oral gavaged every 4 days for 28 days with TCDD (n=3). **(E)** Circadian regulated genes are denoted with a “Y”. An orange ‘X’ indicates abolished diurnal rhythm following oral gavage with 30 μg/kg TCDD every 4 days for 28 days. ZT indicates statistically significant (P1(*t*) > 0.8) time of maximum gene induction/repression. Counts represent the maximum number of raw aligned reads for any treatment group. Low counts (<500 reads) are denoted in yellow with high counts (>10,000) in pink. Differential expression with a posterior probability (P1(*t*)) >0.80 is indicated with a black triangle in the top right tile corner. Protein subcellular locations were obtained from COMPARTMENTS and abbreviated as: cytosol (C), mitochondrion (M), mitochondrial outer membrane (OMM), mitochondrial inner membrane (IMM), nucleus (N), peroxisome (P), and plasma membrane (PM). **(F/H)** Capillary electrophoresis was used to assess ACOX1 and ECHS1 protein levels in total lysate prepared from liver samples harvested between ZT0-3 (n=3). Bar graphs denote the mean ± SEM. Statistical significance (^*^p ≤ 0.05) was determined using a one-way ANOVA followed by Dunnett’s post-hoc analysis. **(G)** Diurnal expression of *Acox1* was assessed by RNA-seq (n=3). Asterisks denotes a posterior probability (P1(*t*) > 0.8) within the same timepoint comparing vehicle to TCDD. Diurnal rhythmicity (‡) was determined using JTK_CYCLE for each treatment. Circadian data are plotted twice along the x-axis to better visualize the gene expression rhythmicity. **(I)** ECHS1 activity was assessed by monitoring the depletion of crotonyl-CoA which has an absorbance at 263 nm. Statistical significance (^*^p ≤ 0.05) was determined using a one-way ANOVA followed by Dunnett’s post-hoc analysis.

### Hydration of trans-2-enoyl-CoA

The hydration of trans-2-enoyl-CoA to 3-hydroxyacyl-CoA is catalyzed by enoyl-CoA hydratases (**Fig. 5**). In the mitochondria, trifunctional protein (MTP) encoded by *Hadha* or *Echs1* and *Hadhb*, preferentially hydrates, oxidizes, and cleaves the resulting 3-ketoacyl-CoA to produce acetyl-CoA and an acyl-CoA that is two carbons shorter. HADHA exhibits the highest activity for C12-16 enoyl-CoAs with virtually no activity towards C4 enoyl-CoAs.^54^ The expression of *Hadha*, which encodes for enoyl-CoA hydratase in MTP, was not affected by TCDD. ECHS1, the short chain enoyl-CoA hydratase, prefers C4 enoyl-CoAs with diminishing activity towards C10 enoyl-CoAs.^55^ Although *Echs1* transcript levels was repressed 2.0-fold in the presence of AHR enrichment and pDREs, protein levels were induced (Fig. 5H). Furthermore, Fig. 5I shows that a 10-fold higher concentration of octenoyl-CoA blocked the hydration of crotonyl-CoA suggesting accumulating octenoyl-CoA levels inhibit ECHS1 activity.

In peroxisomes, *Ehhadh*, which encodes for the enoyl-CoA hydratase subunit of the liver bifunctional enzyme (L-BPE), was induced 7.7-fold in the absence of AHR enrichment and pDREs. After several cycles, the resulting peroxisomal medium-chain acyl-CoAs are transported to the mitochondria for further β-oxidation. Despite the expression of multiple acyl-CoA dehydrogenases that produce medium-chain enoyl-CoAs, there is only one enoyl-CoA hydratase, ECHS1, with a preference for medium and short-chain acyl-CoAs. Consequently, *Echs1* repression may contribute to the dose-dependent accumulation of octenoyl-CoA (**Table 1, Fig. 5**).

*Eci1* and *2*, and *Decr1* and *2* isomerize mono- and poly-unsaturated acyl-CoA *cis* double bonds to the 2-*trans* configuration, and the reduction of 2,4-dienoyl-CoAs to trans-3-enoyl CoA, respectively (**Supp Fig. 1**). *Eci1, Eci2 and Decr2* were repressed by TCDD (2.1-, 2.2-, and 2.9-fold, respectfully). All four genes exhibited AHR enrichment in the presence of a pDRE. Without appropriate standards, it was not possible to determine the presence of *cis*-versus *trans*-enoyl-CoA or 2-versus 3-enoyl-CoA.

### Dehydrogenation of 3-hydroxyacyl-CoA

The oxidation of 3-hydroxyacyl-CoA to 3-ketoacyl-CoA is catalyzed by 3-hydroxyacyl-CoA dehydrogenase (**Fig. 5**). In the mitochondria, long-chain 3-hydroxyacyl-CoAs are oxidized by the highly expressed alpha subunit (*Hadha*) of MTP which was not affected by TCDD. In peroxisomes, the highly expressed *Ehhadh* subunit, which was induced 7.7-fold, preferentially oxidizes very long- and long-chain 3-hydroxyacyl-CoAs. Medium- and short-chain 3-hydroxyacyl-CoAs are then oxidized by *Hadh* and *Hsd17b4* which were repressed 1.5- and 1.8-fold, respectively. *Hadh* and *Hsd17b4* both exhibited AHR enrichment in the presence of a pDRE while *Ehhadh* induction occurred in the absence of AHR enrichment or pDREs.

### Thiolysis of 3-ketoacyl-CoA

Thiolytic cleavage of 3-ketoacyl-CoA requires an additional CoASH to generate acetyl-CoA and an acyl-CoA that is two carbons shorter (**Fig. 5**). In the mitochondria, the MTP beta subunit (*Hadhb*), which exhibited no change, catalyzes the cleavage of long-chain 3-ketoacyl-CoAs. In peroxisomes, ACAA1A (*Acaa1a*), ACAA1B (*Acaa1b*) and SCP2 (*Scp2*), which preferentially catalyze very long 3-ketoacyl-CoA thiolysis, were repressed 1.8-, 3.9- and 3.5-fold, respectively. Long-, medium- and short-chain 3-ketoacyl-CoAs undergo thiolytic cleavage catalyzed by ACAA2 (*Acaa2*) and ACAT1 (*Acat1*) which were repressed 4.2- and 3.0- fold. All six genes with thiolytic cleavage activity exhibited AHR enrichment in the presence of a pDRE. Collectively, 5 of 6 genes associated with thiolase activity were repressed by TCDD.

### Acyl-CoA Deactivation

Excess FAs can overload β-oxidation causing mitochondrial stress that reduces flux and depletes free CoASH needed for other metabolic pathways including the TCA cycle and FA metabolism. In response, acyl-CoA thioesterases (ACOTs) hydrolyze acyl-CoAs releasing the CoASH from activated FAs (**Fig. 6**).^56^ Therefore, we examined the effect of TCDD on cytosolic and mitochondrial thioesterase expression. *Acot1* and *Acot2*, which exhibit a preference for long-chain acyl-CoAs, were induced 8.9- and 21.2-fold, respectively. ACOT2 is the primary mitochondrial thioesterase with little activity for <10C acyl-CoAs. In contrast, *Acot7* and *Acot13*, are enzymatically inhibited by CoASH, and were repressed 2.4- and 2.8-fold, respectively. Peroxisomal thioesterase *Acot4*, which prefers very long- and long-chain acyl-CoAs was induced 3.7-fold, while CoASH sensitive *Acot8* that exhibits broad substrate preferences (C2-20 acyl-CoAs), was repressed 1.5-fold. Unlike the mitochondria, which catalyzes complete FA oxidation, peroxisomes only complete 2-5 β-oxidative cycles producing medium-chain acyl-CoAs that are hydrolyzed by ACOT3 (*Acot3*; induced 12.5-fold) and readily taken up by the mitochondria. Alternatively, they are transported to the mitochondria following conversion to medium-chain acylcarnitine by carnitine O-octanoyltransferase (*Crot*) (**Fig. 7**). Mitochondrial *Acot9*, which preferentially hydrolyzes medium- and short-chain acyl-CoAs, and is inhibited by CoASH, was also induced 6.0-fold (**Fig. 6**). Accumulating mitochondrial CoASH could then either be used to reactivate free FAs in the matrix or be exported to the cytosol via PMP34 (*Slc25a17*, no expression change) initiating a futile cycle (**Fig. 7**).^56^ *Nudt7* which preferentially cleaves medium-chain acyl-CoAs to acyl-phosphopantetheines and 3’,5’-ADP, was repressed 8.0-fold.^57^ This futile cycle may be exacerbated by the 23.3-fold induction of *Ucp2* which would not only facilitate the mitochondrial export of liberated FAs and FA peroxides, but dissipate the proton gradient and uncouple oxidative phosphorylation.^58,59^ In summary, TCDD-elicited differential gene expression of thioesterases favors the hydrolysis of very long- and long-chain acyl-CoAs freeing CoASH required for other reactions while leaving medium-chain acyl-CoAs intact for peroxisomal and mitochondrial β-oxidation.

**Fig 6:**
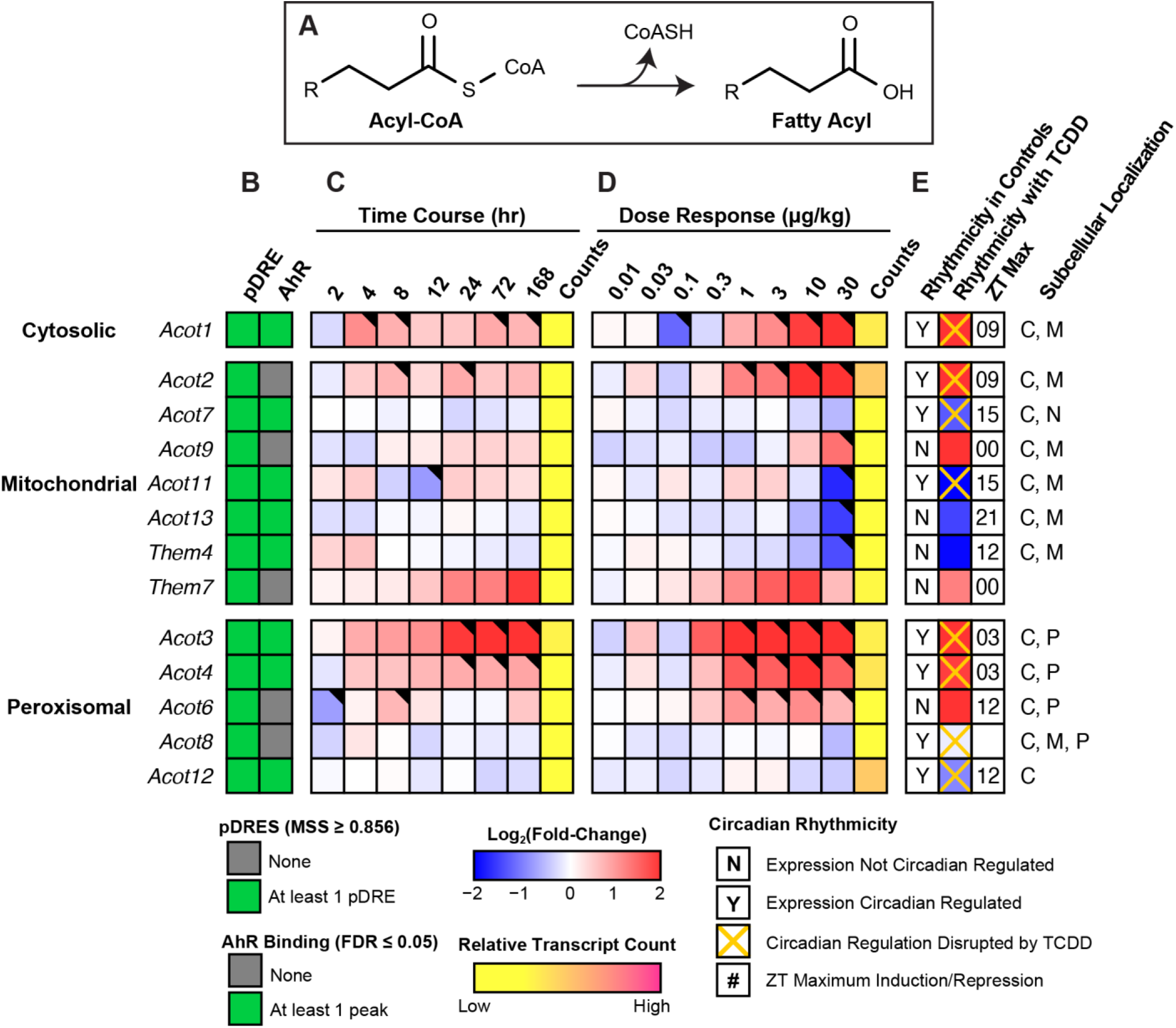
Hydrolysis of acyl-CoAs. Differential expression of genes associated with fatty acid deactivation assessed by RNA-seq. **(A)** Fatty acid deactivation reaction catalyzed via acyl-CoA thioesterases. Official gene symbol designated in the MGI database are listed. **(B)** The presence of putative dioxin response elements (pDREs) and AHR enrichment at 2 hrs. (**C)** Time-dependent gene expression was assessed following a single bolus dose of 30 μg/kg TCDD (n=3). **(D)** Dose-dependent gene expression following oral gavaged every 4 days for 28 days with 0.01, 0.03, 0.1, 0.3, 1, 3, 10 or 30 μg/kg TCDD (n=3). **(E)** Circadian regulated genes are denoted with a “Y”. An orange ‘X’ indicates abolished diurnal rhythm following oral gavage with 30 μg/kg TCDD every 4 days for 28 days. ZT indicates statistically significant (P1(*t*) > 0.8) time of maximum gene induction/repression. Counts represents the maximum number of raw aligned reads for any treatment group. Low counts (<500 reads) are denoted in yellow with high counts (>10,000) in pink. Differential expression with a posterior probability (P1(*t*)) >0.80 is indicated with a black triangle in the top right tile corner. Protein subcellular locations were obtained from COMPARTMENTS and abbreviated as: cytosol (C), mitochondrion (M), nucleus (N), and peroxisome (P).

**Fig 7:**
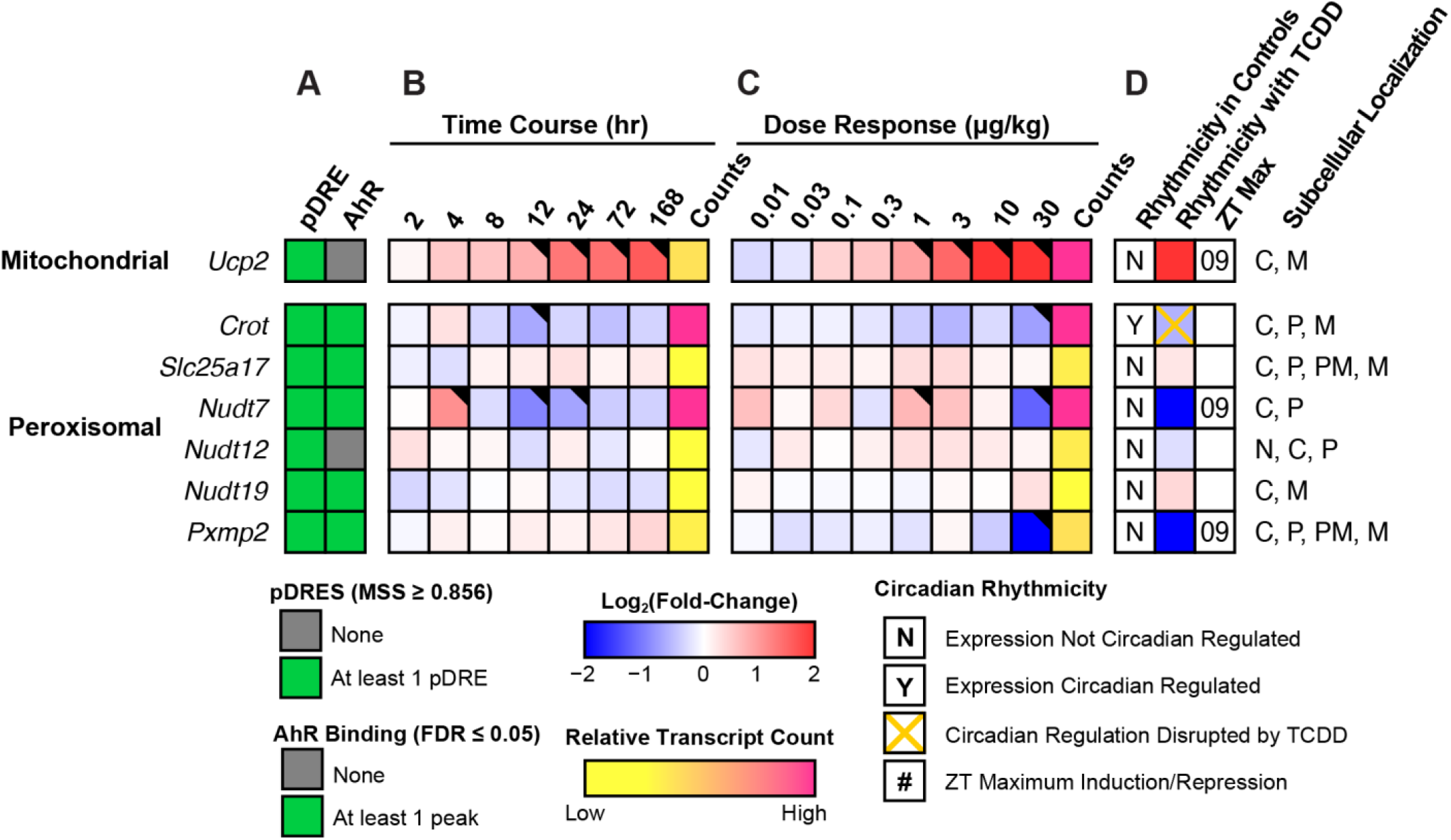
Mitochondria and peroxisomal fatty acid and CoASH transport. Differential expression of genes associated with mitochondrial and peroxisomal fatty acid and CoASH transport assessed by RNA-seq. Mouse Genome Informatics (MGI) official gene symbols are used. **(A)** The presence of putative dioxin response elements (pDREs) and AHR genomic binding at 2 hrs. **(B)** Time-dependent expression following a single bolus dose of 30 μg/kg TCDD (n=3). **(C)** Dose-dependent gene expression assessed following oral gavage every 4 days for 28 days with TCDD (n=3). **(D)** Diurnal regulated gene expression denoted with a “Y”. An orange ‘X’ indicates abolished diurnal rhythm following oral gavage with 30 μg/kg TCDD every 4 days for 28 days. ZT indicates statistically significant (P1(*t*) > 0.8) time of maximum gene induction/repression. Counts represent the maximum number of raw aligned reads for any treatment group. Low counts (<500 reads) are denoted in yellow with high counts (>10,000) in pink. Differential gene expression with a posterior probability (P1(*t*)) >0.80 is indicated by a black triangle in the top right tile corner. Protein subcellular locations were obtained from COMPARTMENTS and abbreviated as: cytosol (C), mitochondrion (M), nucleus (N), peroxisome (P), and plasma membrane (PM).

### Omega(ω)-oxidation of fatty acids

FAs can also undergo ω-hydroxylation with subsequent metabolism by alcohol and aldehyde dehydrogenases to produce dicarboxylic acids (DCAs) when β-oxidation is impeded. Consequently, we examined our metabolomics data for the presence of DCAs. Dose-dependent changes in short-, medium- and long-chain DCAs were identified (**Table 2**). MCFAs (C10:0 and C12:0) are reported to have the greatest affinity for ω-oxidation.^60^ Sebacic (C10:0) and dodecanedioic (C12:0) acids were induced 20.4- and 1.5-fold. Both can undergo further peroxisomal β-oxidation to produce shorter DCAs. Suberic (C8:0), azelaic (C9:0) and undecanedioic (C11:0) acids were induced 2.8-, 2.7-, and 8.9-fold, respectively, between 1 and 10 µg/kg TCDD with decreasing levels at higher doses. The detection of azelaic acid has been previously reported in TCDD treated mice.^37^

**Table 2:** Induction of dicarboxylic acid levels by TCDD. Dicarboxylic acid levels were assessed in mice using untargeted liquid chromatography tandem mass spectrometry. Mice (n=4-5) were orally gavaged every 4 days for 28 days with sesame oil vehicle or TCDD. Bold font and asterisks (^*^) denote statistical significance (*p* ≤ 0.05) determined using a one-way ANOVA with Dunnett’s *post-hoc* analysis. Scores were determined by Progenesis with 60 being the maximum value and 0 being the minimum value. Scores ranging from 30 − 40 are based on mass error and isotope distribution similarity, while score >40 are based on mass error, isotope distribution and fragmentation score. All identified compounds have a score distribution averaging ∼35.

| Compound ID | Description | C<sub>n</sub> | Score | Fold-Change (TCDD vs. Veh.) |  |  |  |  |
| --- | --- | --- | --- | --- | --- | --- | --- | --- |
|  |  |  |  | 0.3 µg/kg | 1 µg/kg | 3 µg/kg | 10 µg/kg | 30 µg/kg |
| HMDB00672 | Hexadecanedioic acid | 16 | 46.1 | 1.00 ± 0.03 | 0.93 ± 0.04 | 1.10 ± 0.14 | 1.33 ± 0.17 | **1.90 ± 0.41\*** |
| HMDB00872 | Tetradecanedioic acid | 14 | 43.5 | 0.99 ± 0.02 | 1.10 ± 0.07 | 0.91 ± 0.11 | **0.38 ± 0.04\*** | **0.57 ± 0.10\*** |
| HMDB00623 | Dodecanedioic acid | 12 | 37.9 | 0.95 ± 0.01 | 1.13 ± 0.05 | 1.12 ± 0.05 | 1.18 ± 0.07 | **1.53 ± 0.11\*** |
| HMDB00888 | Undecanedioic acid | 11 | 42.7 | 0.36 ± 0.02 | 2.05 ± 0.21 | 3.01 ± 1.38 | **8.79 ± 0.67\*** | **5.40 ± 1.96\*** |
| HMDB00792 | Sebacic acid | 10 | 36.5 | 0.44 ± 0.06 | 3.67 ± 0.53 | 6.54 ± 3.33 | **20.39 ± 3.53\*** | **12.79 ± 5.62\*** |
| HMDB00784 | Azelaic acid | 9 | 45.9 | 0.90 ± 0.09 | **2.73 ± 0.26\*** | **2.32 ± 0.57\*** | 1.66 ± 0.12 | **2.22 ± 0.20\*** |
| HMDB00893 | Suberic acid | 8 | 37.9 | 0.83 ± 0.02 | 1.54 ± 0.13 | 1.50 ± 0.32 | **2.82 ± 0.38\*** | **2.06 ± 0.31\*** |

In ω-oxidation, SCFAs, MCFAs, LCFAs and VLCFAs are first ω-hydroxylated by the CYP4Bs, CYP4As, CYP4Fs or CYP4Us. However, most cytochrome P450, ADH, and ALDH genes associated with ω-oxidation were repressed (**Fig. 8**). Specifically, *Cyp4a12a, Cyp4a12b* and *Cyp4a32* which preferentially ω-hydroxylate MCFAs were repressed 150.9-, 23.6 and 1.9-fold, respectively. Similarly, *Cyp4f13* (−3.2-fold), *Cyp4f14* (−387.6-fold), *Cyp4f15* (−12.2-fold), *Cyp4f17* (−3.1-fold) and *Cyp4v3* (−4.5-fold) that preferentially ω-hydroxylate LCFAs were all repressed. *Cyp2u1* which mediates the ω-hydroxylation of LCFAs was also repressed 10.2-fold.

**Fig 8:**
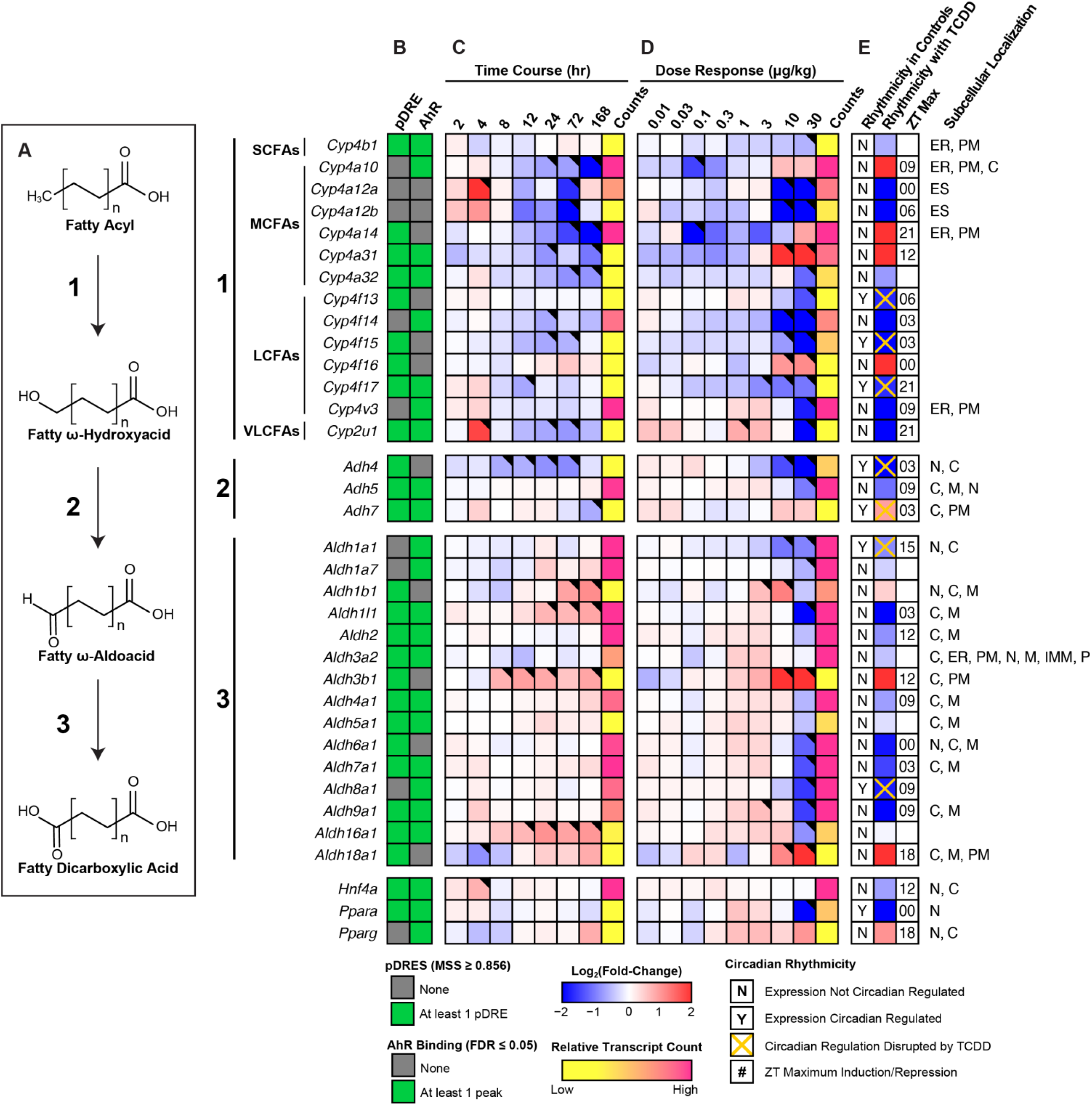
ω-Oxidation of fatty acids into dicarboxylic acids. Differential expression of genes associated with ω-oxidation assessed by RNA-seq. **(A)** Fatty acid ω-oxidation is sequentially catalyzed via CYP450s, alcohol dehydrogenases and aldehyde dehydrogenases. Official gene symbol designated in the MGI database are listed. **(B)** The presence of putative dioxin response elements (pDREs) and AHR enrichment at 2 hrs. **(C)** Time-dependent gene expression was assessed following a single bolus dose of 30 μg/kg TCDD (n=3). **(D)** Dose-dependent gene expression following oral gavaged every 4 days for 28 days with TCDD (n=3). **(E)** Circadian regulated genes are denoted with a “Y”. An orange ‘X’ indicates abolished diurnal rhythm following oral gavage with 30 μg/kg TCDD every 4 days for 28 days. ZT indicates statistically significant (P1(*t*) > 0.8) time of maximum gene induction/repression. Counts represent the maximum number of raw aligned reads for any treatment group. Low counts (<500 reads) are denoted in yellow with high counts (>10,000) in pink. Differential expression with a posterior probability (P1(*t*)) >0.80 is indicated with a black triangle in the top right tile corner. Protein subcellular locations were obtained from COMPARTMENTS and abbreviated as: cytosol (C), endoplasmic reticulum (ER), extracellular space (ES), mitochondrion (M), inner mitochondrial membrane (IMM), nucleus (N), peroxisome (P), and plasma membrane (PM).

**Fig 9:**
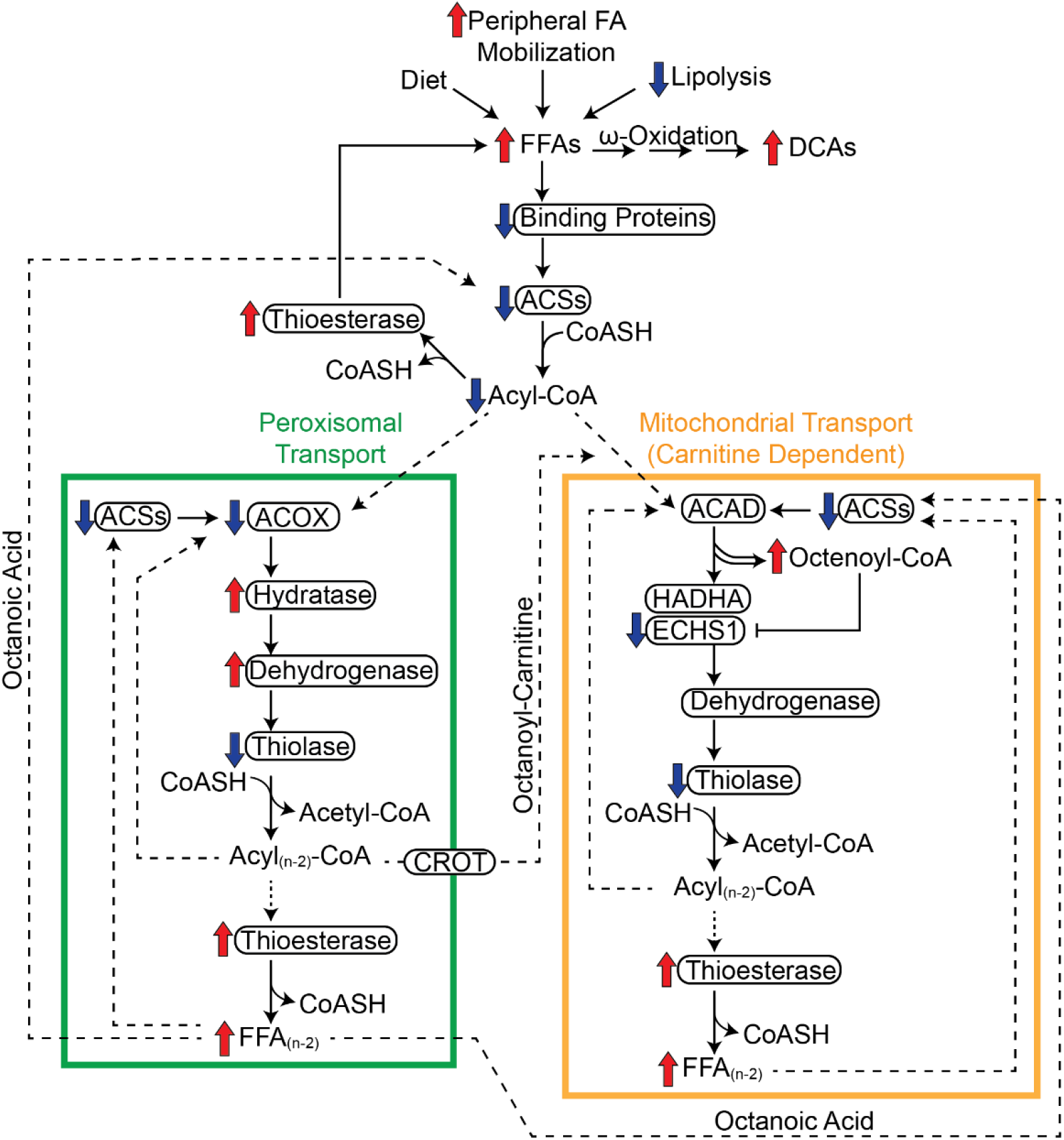
Summary of the effects of TCDD on fatty acid oxidation. Inhibition of β-oxidation in TCDD-induced hepatic steatosis involves the (i) repression of FA binding proteins, (ii) repression of FA activation, and (iii) induction of thioesterase activity. Acyl-CoA synthetase repression reduces activated FAs available for β-oxidation while the repression of binding proteins increases acyl-CoA susceptibility to hydrolysis by induced thioesterases. VLCFAs and LCFAs were partially metabolized in peroxisomal β-oxidation (green box) to medium-length chain FAs (MCFAs) due to peroxisomal specific thioesterase activity which deactivates acyl-CoAs. Peroxisomal CROT transports MCFAs out of peroxisomes to the mitochondria (orange box). MCFAs, such as octanoic acid, could also diffuse out of peroxisomes and into mitochondria where they undergo oxidation following activation by acyl-CoA synthetases. In the canonical pathway, acyl-CoAs are first oxidized by acyl-CoA dehydrogenases that were minimally affected by TCDD. Two enzymes catalyze the subsequent step of mitochondrial β-oxidation; HADHA (which prefers long chain acyl-CoAs) and ECHS1 which prefers short and medium chain acyl-CoAs. Peroxisomal and mitochondrial medium chain acyl-CoA species would accumulate, specifically octenoyl-CoA, due to substrate overload and inhibition of ECHS1 activity by accumulating octenoyl-CoA levels. Octenoyl-CoA accumulation also sequesters free CoASH inhibiting other CoASH-dependent reactions including the thiolytic cleavage of the ketoacyl-CoA into acetyl-CoA and an acyl-CoA that is 2 carbons shorter. Therefore, TCDD dose-dependently induced a futile cycle of FA activation and acyl-CoA hydrolysis leading to octenoyl-CoA accumulation that inhibits ECHS1 activity and depletes free coenzyme A required by enzymes in other pathways leading to the inhibition of β-oxidation and energy depletion. Proteins are denoted with an oval. Red arrows indicate induction and blue arrows indicate repression. Solid lines denote reactions while dashed lines denote substrate trafficking.

However, the principal genes that mediate FA ω-hydroxylation, *Cyp4a10* and *Cyp4a14*, were induced 4.2-, and 6.7-fold, respectively. *Cyp4a10* and *Cyp4a14* induction is primarily mediated by PPARA (*Ppara*), which was repressed 8.7-fold, with AHR enrichment in the presence of a pDRE. *Cyp4a31* and *Cyp4f16* were also induced 25.9- and 4.1-fold. Of these induced genes, only *Cyp4a31* exhibited AHR enrichment with a pDRE by TCDD at 2 hrs. Poorly defined alcohol and aldehyde dehydrogenases are responsible for the oxidation of ω-hydroxylated FAs. *Adh4* and *Adh5* expression was repressed 14.8- and 2.2-fold, respectively, consistent with other reports.^61,62^ TCDD did not affect *Adh7* which is reported to oxidize long ω-hydroxylated FAs.^63^ Highly expressed *Aldh1a1* was repressed 1.9-fold, while *Aldh3a2* showed negligible changes. However, *Aldh3b1* and *Aldh18a1* were induced 41.1- and 19.0-fold, respectively. Despite most genes being repressed, key ω-oxidation genes were induced by TCDD consistent with increased DCA levels.

## DISCUSSION

In this study, TCDD is used as a prototypical AHR ligand and represents the cumulative burden of all AHR agonists. The dose range and regimen approached steady state levels while inducing full dose response curves for many genes in the absence of (i) necrosis or apoptosis, (ii) significant serum ALT increases, (iii) changes in food consumption and (iv) body weight loss >15%.^8,9^ To provide perspective, 30 µg/kg TCDD resulted in mouse hepatic tissue levels comparable to serum levels reported in Viktor Yushchenko following intentional poisoning, while 0.03-0.1 μg/kg resulted in serum levels comparable to the Seveso cohort of women following the 1976 chemical release accident. At 0.01 µg/kg, hepatic levels were comparable to control hepatic levels, and to dioxin-like compound levels in US, German, Spanish and British serum samples.^8,64,65^ Consequently, the dose range and regimen are relevant to human exposures, and the elicited effects cannot be attributed to overt toxicity. Conservation of the AhR, and similarities in AHR-mediated dyslipidemia and metabolic disease between rodents and humans, suggest a common mechanism that may identify novel therapeutic interventions for NAFLD which currently has limited treatment options.^66^

TCDD dose-dependently induced steatosis in mice with marked increases in TAGs, CEs, phospholipids, and free FAs in the absence of acute toxicity.^13,15^ Previous reports of β-oxidation inhibition by TCDD^13,17^ were confirmed by dose-dependent decreases in acyl-CoA levels, consistent with reported decreases of acyl-CoAs by 2,3,7,8-tetrachlorodibenzo-*p*-furan (TCDF) in mice and in human liver Huh-7 cells following TCDD treatment.^67,68^ We further investigated this inhibition by integrating metabolomics, ChIP-seq, and RNA-seq datasets to determine the effects of TCDD on FA oxidation, focusing on differential gene expression associated with lipid hydrolysis, FA activation, binding proteins, β-oxidation, and acyl-CoA hydrolysis.

RNA-seq analysis identified seven of nine lipid hydrolases repressed by TCDD with only *Pnpla2* and *Lpl* showing induction. Yet, TCDD increased hepatic free FA levels which could serve as substrate for β-oxidation. The increase in FA levels is attributed not only to increased hepatic uptake of dietary FAs and mobilized peripheral lipids following CD36 induction, but also reduced ACS levels.^13,15,17^ ACSs catalyzes a two-step, ATP-dependent, reaction producing activated FAs required for β-oxidation to proceed. In addition, genes encoding the predominant FA binding proteins FABP1, DBI, and SCP2, which also bind acyl-CoAs, heme, bile acids, retinoids, and other hydrophobic ligands, were repressed by TCDD, therefore reducing cytosolic, peroxisomal, and mitochondrial acyl-CoA binding capacity and trafficking. Furthermore, FABPs, DBI, and SCP2 buffer the toxicity, block the signaling potential of free ligands, and protect acyl-CoAs from hydrolysis.^31,69^ For example, long-chain acyl-CoAs are powerful disruptors of membrane bilayers, inhibit diverse enzyme activities, and alter ion channel function.^70^ Consequently, decreases in ACS and binding proteins would impede FA metabolism by reducing activated FA levels, and increasing acyl-CoA susceptibility to hydrolysis.

CoASH is an obligate cofactor for >100 different metabolic reactions. It does not passively diffuse across membranes, and therefore distinct cytosolic, peroxisomal, and mitochondrial CoASH and acyl-CoA pools exist. Hereditary or acquired conditions involving CoA ester accumulation, often referred to as CoASH sequestration, toxicity, or redistribution (CASTOR) disease, lead to the accumulation of one or more acyl-CoA species that reduce free CoASH levels causing adverse effects.^71^ Deficiencies in short-, medium-, long-, and very long-chain acyl-CoA dehydrogenase, trifunctional protein, and carnitine shuttle activities are associated with CASTOR disease.^71^ This phenomena is also associated with the toxicity of valproic, salicylic, pivalic, phenylbutyric, and benzoic acids and does not solely arise from free CoASH depletion.^71,72^

In response to CoASH sequestration, thioesterases prevent acyl-CoA levels from reaching toxic levels.^35^ TCDD dose-dependently induced cytosolic, peroxisomal, and mitochondrial thioesterases including *Acot9* which prefers medium-chain acyl-CoAs, but is strongly inhibited by CoASH favoring octanoyl-CoA accumulation. Furthermore, *Nudt7* which hydrolyzes free CoASH and acyl-CoAs to (acyl)phosphopantetheines and 3’,5’-ADP, was repressed.^57^ *Nudt7* repression and thioesterase induction would increase free CoASH levels and de-activate FAs in an ATP-consuming futile cycle that inhibits β-oxidation. Collectively, (i) ACS repression and thioesterase induction would reduce activated FA levels required for β-oxidation, while (ii) decreased FABPs, DBI, and SCP2 binding capacity would increase acyl-CoA susceptibility to hydrolysis and compromise trafficking substrates to metabolic pathways. Moreover, *Ucp2* induction would export liberated FAs from the matrix, uncoupling oxidative phosphorylation and further compromise energy production. The accumulation of TAGs and CEs containing long-chain FAs,^13^ the dose-dependent decrease in acyl-CoA levels (**Table 1**), and the detection of palmitoylcarnitine in serum, all suggest TCDD impaired FA oxidation.^15^ Increased palmitoyl (C16)-, tetradecanoyl (C14)-, and decanoyl (C10)-carnitine levels are also reported in factory workers with 29.49-765.35 pg TEQs/g lipid levels.^73^ Elevated serum acylcarnitine levels are associated with NAFLD,^74^ suggesting the effects of TCDD in mice may have relevance in humans.

An unexpected result was the dose-dependent increase in octenoyl-CoA. In the mitochondria, long-chain acyl-CoAs are oxidized by multiple dehydrogenases that were minimally affected by TCDD. The trifunctional protein (MTP), encoded by *Hadha* or *Echs1* and *Hadhb*, carries out the enoyl-CoA hydratase, hydroxyacyl-CoA dehydrogenase, and 3-ketothiolase activities. *Hadha* expression was not affected by TCDD, *Hadhb* was modestly induced, and thiolases (*Acaa1b, Acaa2*) were repressed. Enoyl-, 3-hydroxy- and 3-keto-acyl-CoA intermediates are not substrates for thioesterases. Very long- and long-chain acyl-CoAs that escape thioesterase hydrolysis and complete several β-oxidation spirals would result in the production of medium-chain acyl-CoAs. TCDD induced thioesterases with a preference for very long- and long-chain acyl-CoAs. *Acot9* which prefers medium-chain acyl-CoAs was also induced but is strongly inhibited by CoASH.^35^ Peroxisomal MCFAs produced by peroxisomal β-oxidation would be ferried into the mitochondrial matrix via carnitine O-octanoyltransferase (*Crot*) which prefers medium-chain acyl-CoAs.^33,57^ MCFA trafficking is not dependent on binding proteins and they tend to be metabolized by hepatic mitochondrial β–oxidation following activation as opposed to being incorporated into TAGs, CEs or phospholipids.^51^ We posit that accumulating peroxisomal and mitochondrial octanoyl-CoAs are oxidized to octenoyl-CoA by ACADM which was largely unaffected by TCDD. Accumulating octenoyl-CoA levels represent a metabolic conundrum since the HADHA enoyl-CoA hydratase subunit prefers substrate chain lengths of C12-16 with virtually no activity towards C4 substrates while the ECHS1 subunit prefers C4 enoyl-CoAs with diminishing binding affinity up to C10. This is somewhat analogous to ACADM deficient mice where octanoyl-CoA oxidation is not compensated by other acyl-CoA dehydrogenases.^75^ Moreover, in addition to being a poor substrate for hydration, octenoyl-CoA also inhibits ECHS1 activity.

When β-oxidation is hindered, FAs can undergo ω-hydroxylation, primarily by CYP4A10 and CYP4A14, with subsequent oxidation by poorly defined cytosolic alcohol and aldehyde dehydrogenases to produce DCAs.^76^ In humans, ω-oxidation is a secondary pathway responsible for 5-10% of FA metabolism under normal conditions. During fasting, starvation, or when β-oxidation is impeded due to an inborn error of metabolism, ω-oxidation is considered a rescue pathway producing DCAs that undergo peroxisomal β-oxidation to produce TCA cycle and gluconeogenesis intermediates.^77^ Conversely, DCAs are also associated with oxidative stress, lipotoxicity and PPAR activation that induces β-oxidation.^78^ However, TCDD repressed *Ppara* expression and inhibits PPARα-mediated gene expression,^79^ thus compromising β–oxidation induction by DCAs.

Long-chain DCAs can be reactivated to the corresponding CoA ester and undergo chain shortening by peroxisomal β–oxidation to produce DCAs of varying length following hydrolysis by thioesterases.^80^ For example, dodecanedioic acid (12C) can be oxidized to sebacic (C10) and suberic acids (C8) while undecanedioic acid (11C) is rarely detected since it is partially oxidized in peroxisomes before further mitochondrial metabolism. Interestingly, TCDD decreased serum azelaic (C9) acid levels due to the repression of CES3 that hydrolyzes monoesters to release azelaic acid.^37^ By extrapolation, we surmise that decreasing DCA levels (**Table 2**) at higher TCDD doses is inversely correlated with the repression of *Ces* genes that exhibit different DCA monoester preferences as reported for other carboxyesterase substrates.^37,81^

In summary, our data suggests β-oxidation inhibition by TCDD is due to the differential expression of genes peripheral to the spiral itself. More specifically, TCDD represses lipid hydrolysis, FA activation, and binding protein expression, while inducing thioesterases. However, these studies were limited to effects on gene expression and selected proteins, and do not consider post-translational modifications that can also regulate enzyme activity. The data are also consistent with TCDD inducing a futile cycle of FA activation by ACSs and de-activation by thioesterases that inhibits complete oxidation of FAs, uncouples oxidative phosphorylation, and depletes ATP levels. Consequently, FAs underwent ω-oxidation producing PPAR ligands in an attempt to induce β-oxidation. However, *Ppara* and *Fabp1*, the encoded proteins, of which, deliver ligands to PPARα, were repressed thwarting efforts to induce β-oxidation. This would not only further deplete energy levels but also increase free FA and DCA levels that likely exacerbate TCDD toxicity. The inhibition of ECHS1 activity and accumulation of mitochondrial octenoyl-CoA may also contribute to toxicity by precipitating metabolic decompensation due to the exhaustion of free CoASH required by enzymes in other pathways. Moreover, octenoyl-CoA may itself be toxic by inappropriately inducing signaling, and/or possess lytic (detergent-like) properties that disrupt membranes. Although these effects are consistent with the development and progression of NAFLD, as well as AHR-mediated hepatotoxicity elicited by TCDD and related compounds, their relevance in human models warrants further investigation. This should include complementary metabolomic analyses of other fractions and compartments such as serum and urine for metabolites associated with NAFLD.

## ACKOWLEDGEMENTS

This project was supported by the National Institute of Environmental Health Sciences Superfund Research Program [NIEHS SRP P42ES004911] to TRZ. TRZ was partially supported by AgBioResearch at Michigan State University. GNC and RRF were supported by NIEHS Multidisciplinary Training in Environmental Toxicology [T32ES007255]. KAF was supported by the Canadian Institutes of Health Research Doctoral Foreign Study Award [DFS-140386].

## CONTRIBUTIONS

GNC, RRF and TRZ designed the project. RRF, KAF, NAZ and RN performed the animal work. GNC, RRF, NAZ, and KAF performed the experiments. RRF developed the LC-MS workflow for untargeted metabolomics. GNC, RRF and NAZ compiled all the data and produced the figures and tables. GNC, RRF and TRZ wrote the manuscript. All authors reviewed the manuscript.

---

**Supplementary Figure 1:**
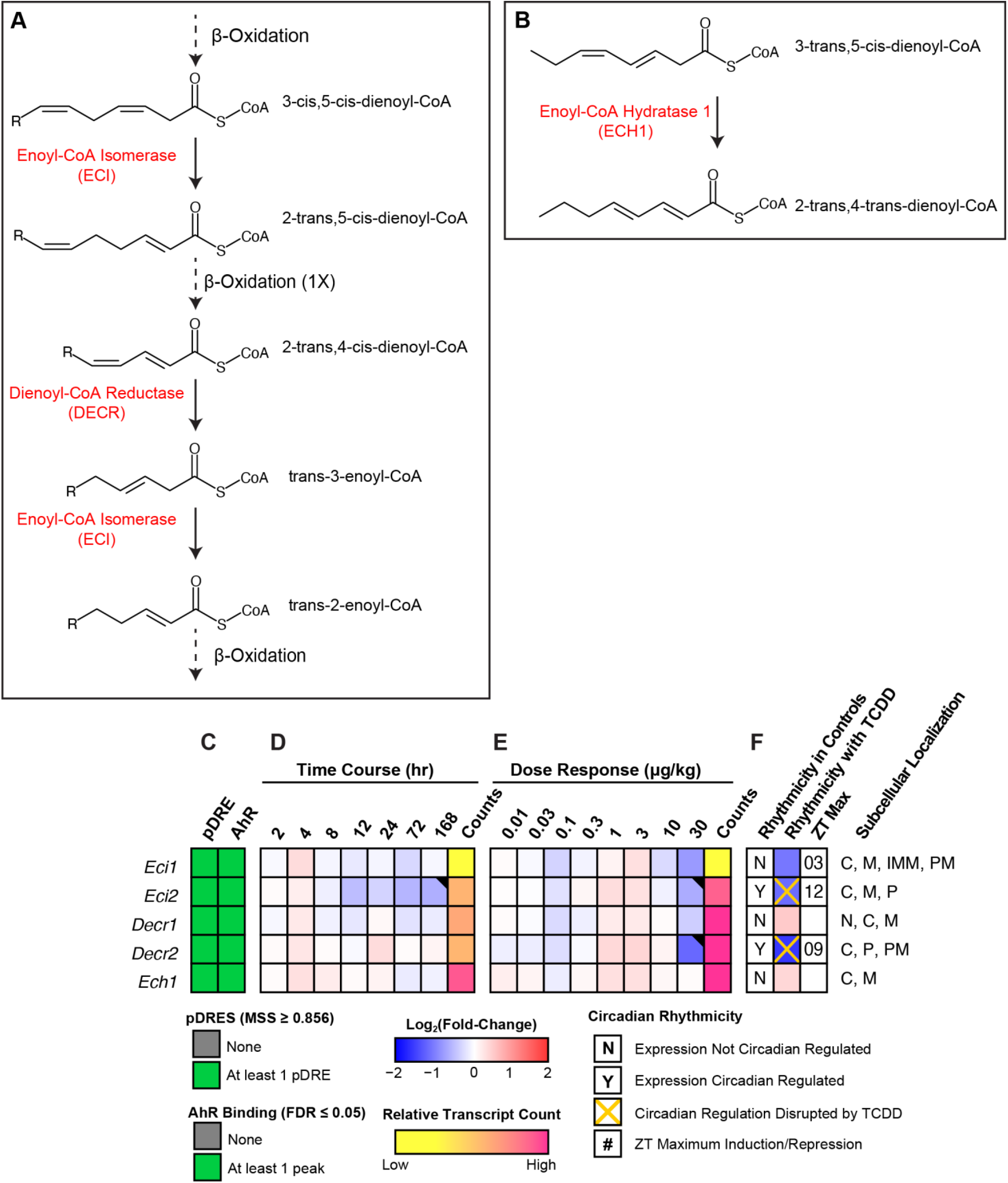
Effect of TCDD on the differential expression of auxiliary genes associated with unsaturated fatty acid oxidation. **(A)** β-Oxidation of polyunsaturated fatty acids involves the use of the two auxiliary enzymes, enoyl-CoA isomerase (ECI) and dienoyl-CoA reductase (DECR). **(B)** β-Oxidation of 3-trans,5-cis-dienoyl-CoA species requires enoyl-CoA hydratase 1 (ECH1) isomerization activity. Official gene symbol designated in the MGI database are listed. Differential expression of genes associated with unsaturated fatty acid metabolism. **(C)** The presence of putative dioxin response elements (pDREs) and AHR enrichment at 2 hrs. **(D)** Time-dependent gene expression was assessed following a single bolus dose of 30 μg/kg TCDD (n=3). **(E)** Dose-dependent gene expression following oral gavaged every 4 days for 28 days with TCDD (n=3). **(F)** Circadian regulated genes are denoted with a “Y”. An orange ‘X’ indicates abolished diurnal rhythm following oral gavage with 30 μg/kg TCDD every 4 days for 28 days. The ZT with maximum gene induction/repression is provided. Counts represent the maximum number of raw aligned reads for any treatment group. Low counts (<500 reads) are denoted in yellow with high counts (>10,000) in pink. Differential expression with a posterior probability (P1(*t*)) >0.80 is indicated with a black triangle in the top right tile corner. Protein subcellular locations were obtained from COMPARTMENTS and abbreviated as: cytosol (C), mitochondrion (M), inner mitochondrial membrane (IMM), peroxisome (P), and plasma membrane (PM).

**Supplementary Table 1:**
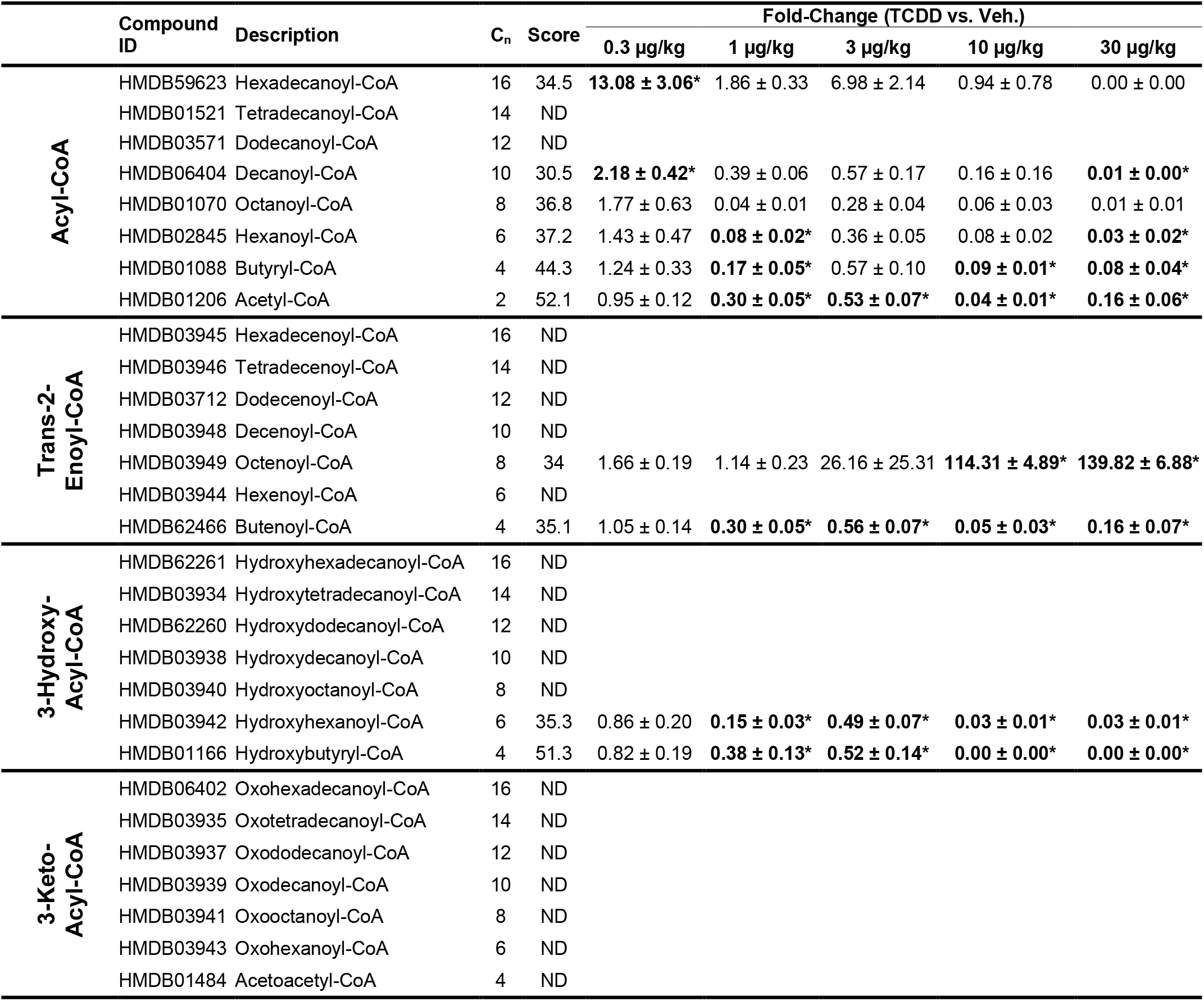
Effect of TCDD on β-oxidation intermediate levels. β-Oxidation intermediate levels were assessed using untargeted liquid chromatography tandem mass spectrometry. Mice (n=4-5) were orally gavaged every 4 days for 28 days with sesame oil vehicle or TCDD. Bold font and asterisks (^*^) denote a statistical significance (*p* ≤ 0.05) as determined using a one-way ANOVA with a Dunnett’s post-hoc analysis. Scores were determined using Progenesis by 60 being the maximum value and 0 being the minimum value. Scores ranging from 30 - 40 are based on mass error and isotope distribution similarity, while score >40 are based on mass error, isotope distribution and fragmentation score. All identified compounds have a score distribution averaging ∼35. Metabolites that were not detected are denoted with “ND”.

